# Post-transcriptional remodelling is temporally deregulated during motor neurogenesis in human ALS models

**DOI:** 10.1101/180372

**Authors:** Raphaelle Luisier, Giulia E. Tyzack, Claire E. Hall, Jernej Ule, Nicholas M. Luscombe, Rickie Patani

## Abstract

Mutations causing amyotrophic lateral sclerosis (ALS) strongly implicate regulators of RNA-processing that are ubiquitously expressed throughout development. To understand the molecular impact of ALS-causing mutations on early neuronal development and disease, we performed transcriptomic analysis of differentiated human control and VCP-mutant induced pluripotent stem cells (iPSCs) during motor neurogenesis. We identify intron retention (IR) as the predominant splicing change affecting early stages of wild-type neural differentiation, targeting key genes involved in the splicing machinery. Importantly, IR occurs prematurely in VCP-mutant cultures compared with control counterparts; these events are also observed in independent RNAseq datasets from SOD1- and FUS-mutant motor neurons (MNs). Together with related effects on 3’UTR length variation, these findings implicate alternative RNA-processing in regulating distinct stages of lineage restriction from iPSCs to MNs, and reveal a temporal deregulation of such processing by ALS mutations. Thus, ALS-causing mutations perturb the same post-transcriptional mechanisms that underlie human motor neurogenesis.

**HIGHLIGHTS:**

- Intron retention is the main mode of alternative splicing in early differentiation.
- The ALS-causing VCP mutation leads to premature intron retention.
- Increased intron retention is seen with multiple ALS-causing mutations.
- Transcriptional programs are unperturbed despite post-transcriptional defects.

**eTOC BLURB:** Luisier et al. identify post-transcriptional changes underlying human motor neurogenesis: extensive variation in 3’ UTR length and intron retention (IR) are the early predominant modes of splicing. The VCP mutation causes IR to occur prematurely during motor neurogenesis and these events are validated in other ALS-causing mutations, SOD1 and FUS.

## INTRODUCTION

Amyotrophic lateral sclerosis (ALS) is a rapidly progressive and incurable condition, which leads to selective degeneration of motor neurons (MNs). ALS-causing mutations implicate crucial regulators of RNA-processing, which are normally expressed throughout development, in the underlying pathogenesis. This raises the hypothesis that post-transcriptional changes occurring at much earlier stages of life, including neurodevelopment, may play a pivotal role in the underlying molecular pathogenesis of ALS. Understanding the impact of ALS-causing mutations on the early stages of neurodevelopment and disease will allow us to elucidate the initiating molecular events. This may guide the development of new therapies targeting primary mechanisms before the disease progresses too far.

Both development and homeostasis of neurons fundamentally rely on the precise implementation of cell-type and stage-specific alternative RNA processing, including alternative splicing (AS) and alternative polyadenylation (APA) (Mansfield and Keene, 2011; Raj et al., 2014; Tian and Manley, 2017; Yap et al., 2016; Zhang et al., 2016). Several mechanisms of AS are established including exon-skipping, mutually exclusive exons and intron retention (IR). Alternative exon usage features particularly in neurons to regulate protein diversity and function (Nilsen and Graveley, 2010). Neural cell-types exhibit a higher proportion of retained introns compared with other tissues and there is an expanding body of evidence demonstrating a functional role for IR both in neuronal development and homeostasis (Braunschweig et al., 2014; Buckley et al., 2011; Mauger et al., 2016; Yap et al., 2012). Increased IR during neuronal differentiation has been shown to down-regulate the expression of transcripts that are unnecessary for mature neuronal physiology (Braunschweig et al., 2014). A recent study showed evidence for post-transcriptional processing of intron-retaining transcripts in response to neuronal activity (Mauger et al., 2016). APA is an alternative mode of RNA-processing that generates distinct 3’ termini most frequently in the 3’ untranslated region (3’ UTR) of mRNA, thereby engendering isoforms of variable 3’ UTR length. 3’ UTRs serve as a key platform in the RNA regulatory network controlling mRNA translational efficiency, localisation and stability (Fabian et al., 2010; Gupta et al., 2014; Will et al., 2013). 3' UTRs harbour extensive tissue-specific length variability that significantly affect their function (Derti et al., 2012). Of all human cell-types, brain-specific isoforms have the longest 3' UTRs (Miura et al., 2013; Wang and Yi, 2014; Zhang et al., 2005). Moreover, mutations perturbing the mRNA secondary structure and sites of mRNA-miRNA interactions can lead to neurodegeneration (Ward and Kellis, 2012). In spite of these findings, the role of 3' UTRs regulation and IR in the context of MN development and disease has remained understudied compared to other forms of alternative splicing.

AS and APA are coordinated by the action of individual or combinations of trans-acting RNA binding proteins (RBPs) that bind to specific sites within (pre-) mRNA (Gabut et al., 2008; Licatalosi et al., 2008). RBPs mediate mRNA splicing, nuclear export and localization. Accumulating evidence implicates RBPs as key regulators of both neurodevelopment and specific forms of neurodegeneration. Indeed, mutations in several RBPs including TDP-43, FUS and TAF15 have been causally linked to familial form of ALS leading to RNA-processing defects in mouse models (Kapeli et al., 2017; Qiu et al., 2014).

However, given that these RBPs are ubiquitously expressed during development, we hypothesized that aberrant splicing and APA play important roles in both development and disease of the nervous system. Against this background we sought to understand how different modes of AS and APA feature in human motor neurogenesis and to examine systematically the impact of ALS-causing mutations on post-transcriptional remodeling during lineage restriction. By combining directed differentiation of induced pluripotent stem cells (iPSCs) with time-resolved RNA sequencing, we first identified the sequential programme of post-transcriptional remodeling that underlies human motor neurogenesis. This allowed us to compare VCP-mutant (VCP*^mu^*) to control cultures during MN differentiation and identify mutation-dependent splicing deregulation in a developmental stage-specific manner. Our findings uncover early post-transcriptional perturbations during motor neurogenesis in ALS, which may induce a state of compensated dysfunction that later impacts motor neuronal integrity, contributing to disease onset and/or progression.

## RESULTS

### Intron retention is the predominant mode of splicing in early motor neurogenesis

To examine post-transcriptional changes during human motor neurogenesis, we performed high-throughput RNA-sequencing (RNA-seq) analyses of polyadenylated RNA isolated from induced pluripotent stem cells (iPSC; day 0), neural precursors (NPC; day 7), ‘patterned’ precursor motor neurons (ventral spinal cord; pMN; day 14), post-mitotic but electrophysiologically immature motor neurons (MN; day 21), and electrophysiologically mature MNs (mMNs; day 35) derived from two patients with the ALS-causing VCP gene mutation and two healthy controls (Figure 1A; 31 samples from 5 time-points and 3 genotypes; 2 clones from 2 healthy controls and 3 clones from 2 ALS patients with VCP mutations: R155C and R191Q). Cellular samples from each stage of MN differentiation have been extensively characterised as previously reported (Hall et al., 2017). Using a set of 19 key gene markers of spinal MN maturation and embryonic development we further confirmed a prior finding that iPSC-derived mMNs resemble fetal rather than adult MNs (Ho et al., 2016) (Figures S1A and S1B). Unsupervised hierarchical clustering (Spearman rank correlation and complete clustering) of the 31 samples using 15,989 reliably expressed genes segregated samples based on their developmental stage within the MN lineage rather than by mutant or control genetic background (Figure 1B).

**Figure 1.**
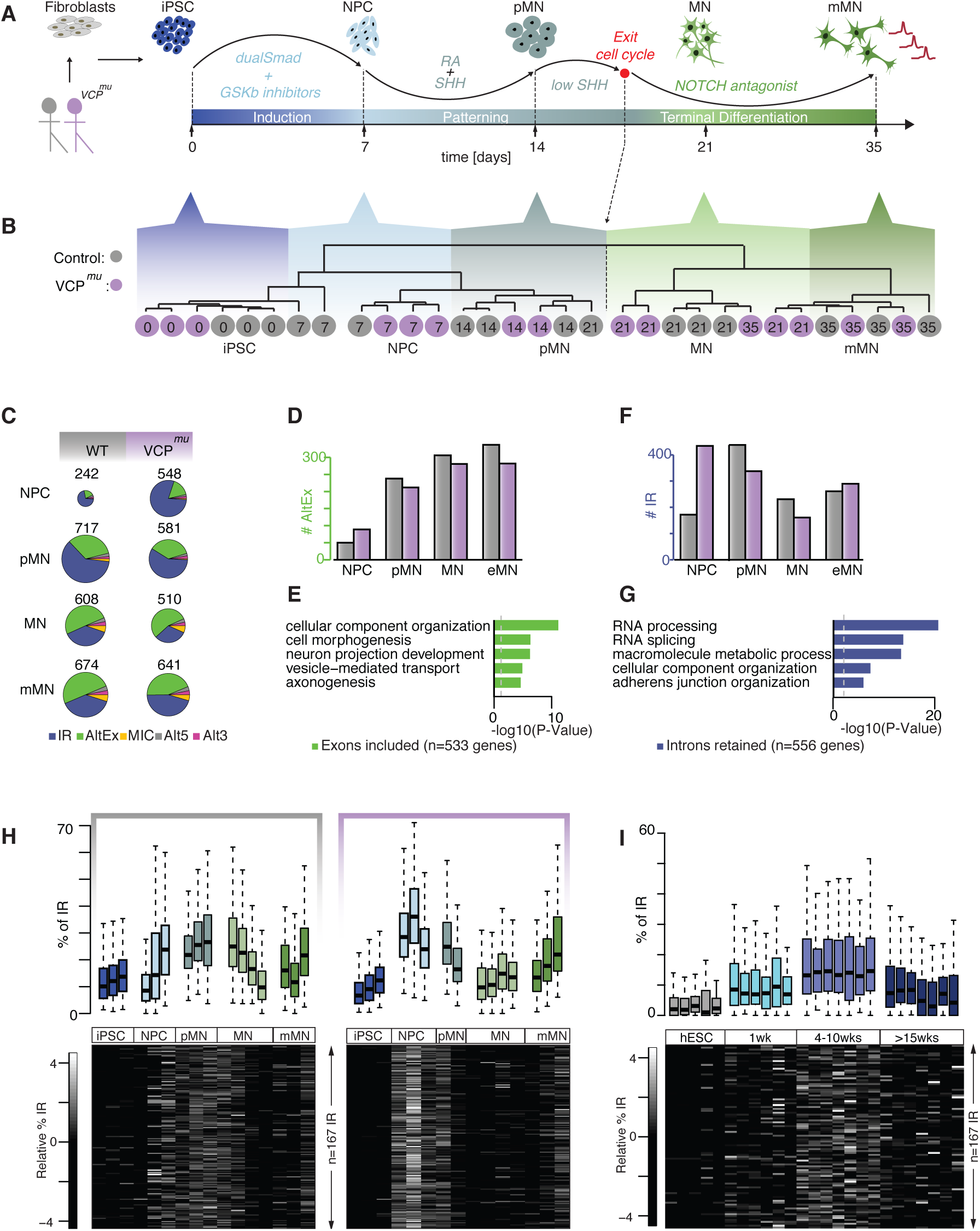
Intron retention is the predominant splicing change at an early stage of MN lineage restriction and occurs prematurely in VCP*^mu^* cultures. **(A)** Schematic showing differentiation strategy for motor neurogenesis. Arrows indicate sampling time-points in days. iPSC clones were obtained from two patients with confirmed VCP mutations (R155C and R191Q; total = 3 different hiPSC lines) and one clone from each of two healthy controls (total = 2 different hiPSC lines). Induced pluripotent stem cells (iPSC); neural precursors (NPC); ‘patterned’ precursor motor neurons (ventral spinal cord; pMN); post-mitotic but electrophysiologically immature motor neurons (MN); electrophysiologically mature MNs (mMN) **(B)** Unsupervised hierarchical clustering of 15,989 genes in n = 31 samples. Grey circles indicate control samples and magenta circles indicate VCP samples; sampling time-points are indicated inside the circles. **(C)** Pie charts representing distributions of regulated splicing events in control and VCP*^mu^* samples at distinct stages of motor neurogenesis compared with the iPSC stage or previous time-point. Intron retention (IR); alternative exon (AltEx); micro-exons (MIC); alternative 5’ and 3’ UTR (Alt5 and Alt3). **(D** and **F)** Bar graphs representing the numbers of exonic and intronic splicing events respectively in controls (grey bars) and VCP*^mu^* samples (magenta bars) at specific time points during motor neuron differentiation. **(E** and **G)** Bar graphs showing the enrichment score of the GO biological pathways associated with transcripts undergoing exonic and intronic splicing events in control samples. **(H)** *(Upper)* Percentage retention for 167 manually curated introns in individual samples at distinct stages of differentiation in healthy controls (*left*) and VCP samples (*right*). Data shown as box plots where the center line is the median, limits are the interquartile range and whiskers are the minimum and maximum. *(Lower)* Heatmap of the standardized relative percentage of IR in individual samples. **(I)** As per **H** but for *in vitro* differentiation of hESCs into an initiation stage (1 week); an NPC stage (4-10 weeks) that produces only neurons upon further differentiation; and after >15 weeks, a cell stage which produces both neurons and glial cells (Wu et al., 2010). See also Figure S1.

We next examined the temporal dynamics of alternative splicing (AS) during MN differentiation. Using the RNA-seq pipeline VAST-TOOLS (Irimia et al., 2014) we identified 1,599 and 1,507 AS events over time in control and VCP*^mu^* samples respectively (Figures S1C and S1D). Consistent with previous studies, > 60% of AS events at later stages of MN terminal differentiation were alternative exon and intron removal events (Figures 1C and S1C,D) (Yap et al., 2012; Zhang et al., 2016). We found that control and VCP*^mu^* samples exhibit a similar progressive increase in cassette exon inclusion over time (Figure 1D) in genes involved in cellular component organization and axonogenesis (Figure 1E).

In contrast, IR accounted for >65% of AS events at the earlier phase of lineage restriction from NPC to pMN (Figure 1C). The expression of intron-retaining transcripts increased at the transition from NPC to pMN (i.e. neural patterning) in control samples (Figure 1F). Over 60% of all IR events were common between VCP and control samples (Figure S1E, *upper left*) - these largely involve genes related to RNA-processing and splicing (Figure 1G). Intriguingly, VCP samples exhibited a striking increase in the expression of intron-retaining transcripts at the earlier developmental transition from iPSC to NPCs (i.e. neural induction; Figure 1F). We found that 53% of IR events started upon patterning in control samples; in contrast 72% of the IR events in VCP*^mu^* cultures started upon neural induction (Figure S1E, *right*). Importantly we found that 60% of the events that started upon neural induction in VCP*^mu^* samples matched with those starting later between induction and patterning in control samples. The remaining events were either unique to VCP*^mu^* cultures or occurred at a different time point. This result indicates that VCP samples exhibit similar IR activity as seen in control counterparts during differentiation, but at a premature stage.

We next conducted Integrative Genomics Viewer (IGV)-guided manual curation to remove low coverage and spurious IR (e.g. some events annotated as IR were found to be alternative 3’UTR after visual inspection). We identified 167 highly confident IR events and assigned a retention value to every intron relative to expression of flanking exons (see Methods). Examining the degree of IR for these selected 167 events across all experimental samples shows a systematically higher percentage of IR in VCP samples at NPC stage compared to control samples of any stage, and confirmed our aforementioned findings of VCP mutation-related premature IR (Figure 1H). We next analyzed our high-confidence set of manually curated IR events in two independent transcriptomic data-sets from human embryonic stem cells (Wu et al., 2010; Yao et al., 2017) confirmed that accumulation of IR during early neurogenesis is a generalizable phenomenon observed across diverse experimental models (Figures 1I and S1F).

VCP plays a key role in the cell cycle and an increased percentage of IR may result from an established link between splicing efficiency and cell cycle (Heyn et al., 2015). We therefore examined the possibility that the apparent acceleration in AS programs observed in VCP*^mu^* samples might be explained by a mutation-dependent increase in cell-cycle activity. In order to address this possibility, we employed flow cytometry on iPSCs, NPCs and pMNs. These experiments effectively excluded differences in cell cycle between VCP*^mu^* and control samples (Figures S1G-I). Collectively, these findings demonstrate that IR is the predominant splicing change that affects early stages of neural lineage restriction. Importantly, IR occurs prematurely in VCP*^mu^* cultures, which cannot be explained by differences in cell-cycle activity.

### A network of splicing regulators exhibit widespread IR in MNs carrying other ALS-causing mutations

We next examined the impact of other ALS-causing mutations by analyzing independent transcriptomic data-sets for familial forms of ALS either caused by mutant SOD1 (n=5; 2 patient-derived SOD1A4V and 3 isogenic control MN samples where the mutation has been corrected) (Kiskinis et al., 2014a) or FUS (n=6; 3 patient-derived FUS R521G and 3 unaffected controls MNs) (Kapeli et al., 2016). We confirmed that IR is the predominant mode of splicing in MNs derived from both mutants (Figure S2A). Importantly, our high-confidence set of 167 IR events affected by premature IR during VCP*^mu^* MN differentiation are also affected in FUS and SOD1 mutant MNs (Figure 2A). 74 and 59 IR events exhibit a significant increase in SOD1 and FUS mutant MNs respectively compared to controls (P-value < 0.01; Fisher count test; Figure S2B). These together form a significantly connected network of proteins enriched in RNA-splicing and translation (Figures 2B and 2C). The extent of splicing for those introns significantly retained in both SOD1*^mu^* and FUS*^mu^* is depicted in a heatmap in Figure 2D. This demonstrates that the premature IR events identified in our ALS model also feature in MNs carrying other ALS-causing mutations. Cumulatively, these findings confirm the generalizability of aberrant IR in different genetic forms of ALS.

**Figure 2.**
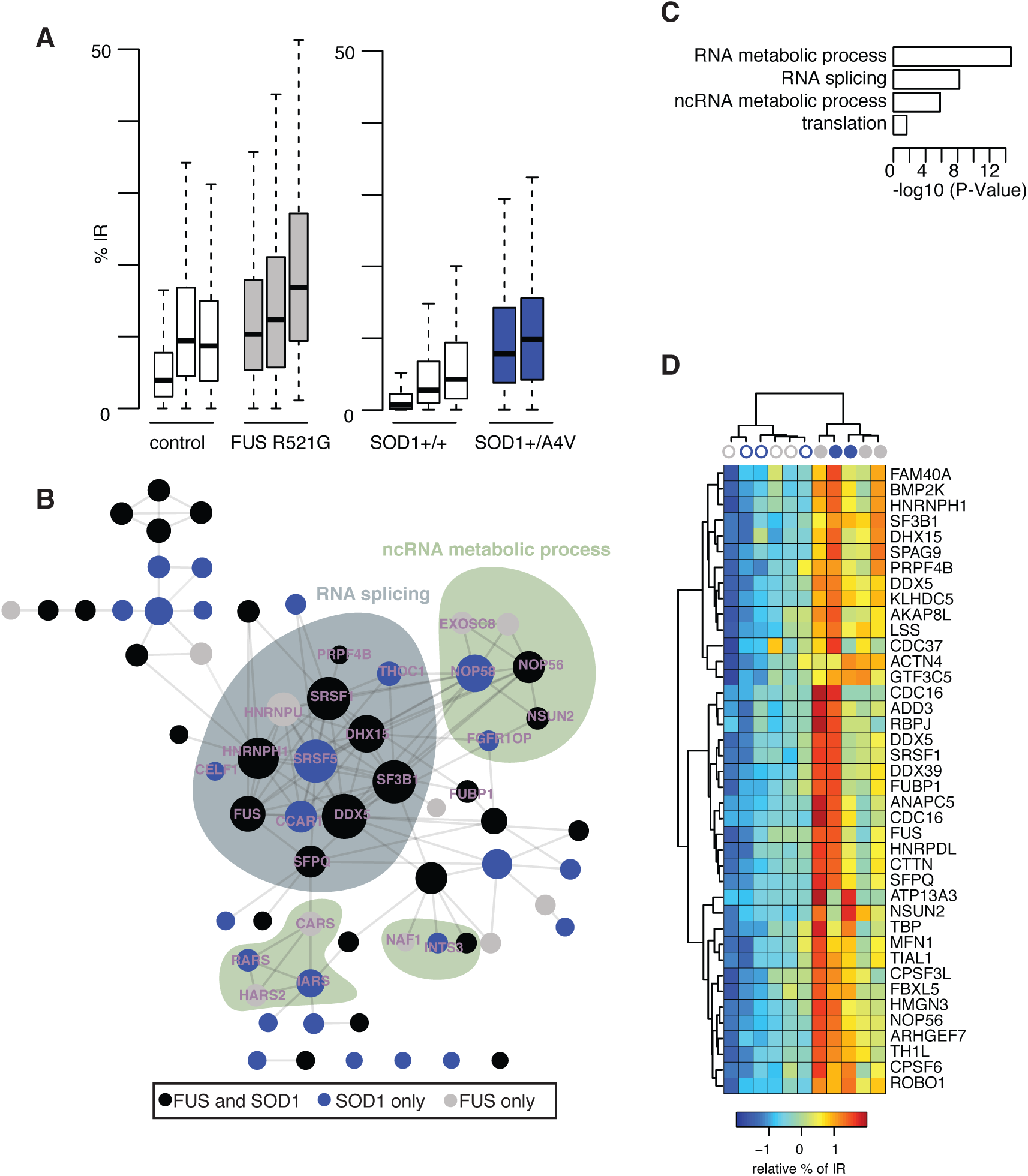
Intron-retaining transcripts involved in neural induction exhibit widespread retention in MN derived from other ALS-causing mutations. **(A)** Percentage retention for 167 manually curated introns in control MNs (white box), FUS R521G mutant MNs (grey box), or SOD1 mutant MNs samples (blue bar) (Kapeli et al., 2016; Kiskinis et al., 2014b). Mutant samples are exhibiting a systematically higher percentage of IR compared with control samples. Data shown as box plot where centre line is the median, limits are the interquartile range and whiskers are the minimum and maximum. **(B)** Network of interacting genes as identified by STRING for the genes that exhibit intron retention during motor neurogenesis and in either SOD1*^mu^* or FUS*^mu^* MNs. Edges represent experimentally determined protein-protein interactions as annotated in STRING data-base (Szklarczyk et al., 2017). The size of the circle is proportional to the number of edges of the gene in the network. **(C)** Selection of GO biological pathways that are enriched among genes that exhibit intron retention during motor neurogenesis and in SOD1*^mu^* or FUS*^mu^* MNs compared with controls. The size of each bar corresponds to the significance -log10(P-value) of the enrichment (P-value from Fisher count test). See also Figure S2. **(D)** Heatmap of the relative intron retention level seen in a selection of 40 genes that exhibit significant intron retention during motor neurogenesis and in both SOD1*^mu^* and FUS*^mu^* MNs. Samples are hierarchically clustered using Manhattan distance and Ward clustering. Intron retention is standardized for each data-set prior analysis. Blue circles for samples from SOD1 study and grey circles for samples from FUS study (empty for control and filled for mutant).

### The most significant IR event identified is in SFPQ, a splicing regulator

The DBHS (Drosophila behavior human splicing) family member SFPQ genes encodes a protein that plays key roles in transcription, splicing, 3’ end processing and axon viability (Cosker et al., 2016; Danckwardt et al., 2007; Dong et al., 2007; Hall-Pogar et al., 2007; Hirose et al., 2014; Patton et al., 1993; Yarosh et al., 2015). SFPQ represents the most significant premature IR event identified during motor neurogenesis in our VCP*^mu^* cultures and was also found to be highly statistically significant in SOD1 and FUS MNs. The SFPQ intron 8/9, ∼9 kb in length, was highly retained during neural patterning in control samples, prematurely (during neural induction) in VCP samples, and in MNs harbouring mutations in FUS and SOD1 (Figures 3A and **3B**). We validated SFPQ intron retention and its temporal deregulation by the VCP-mutation by qPCR analysis of RNA isolated from multiple independent iPSC lines (3 clones from 3 healthy controls and 4 clones from 2 patients with VCP mutations) (Figure 3C and S3A). The key splicing regulators FUS and DDX39 are similarly strongly affected during early lineage restriction by VCP mutation, and by SOD1 and FUS mutations (Figures 3D-I). Changes in IR event were not accompanied by any significant difference in gene expression levels between VCP*^mu^* cultures and controls (Figure S3B). From these results we conclude that distinct ALS-causing gene mutations lead to similar IR events in genes known to have pivotal roles in motor neurogenesis and ALS (Thomas-Jinu et al., 2017; Vance et al., 2009).

**Figure 3.**
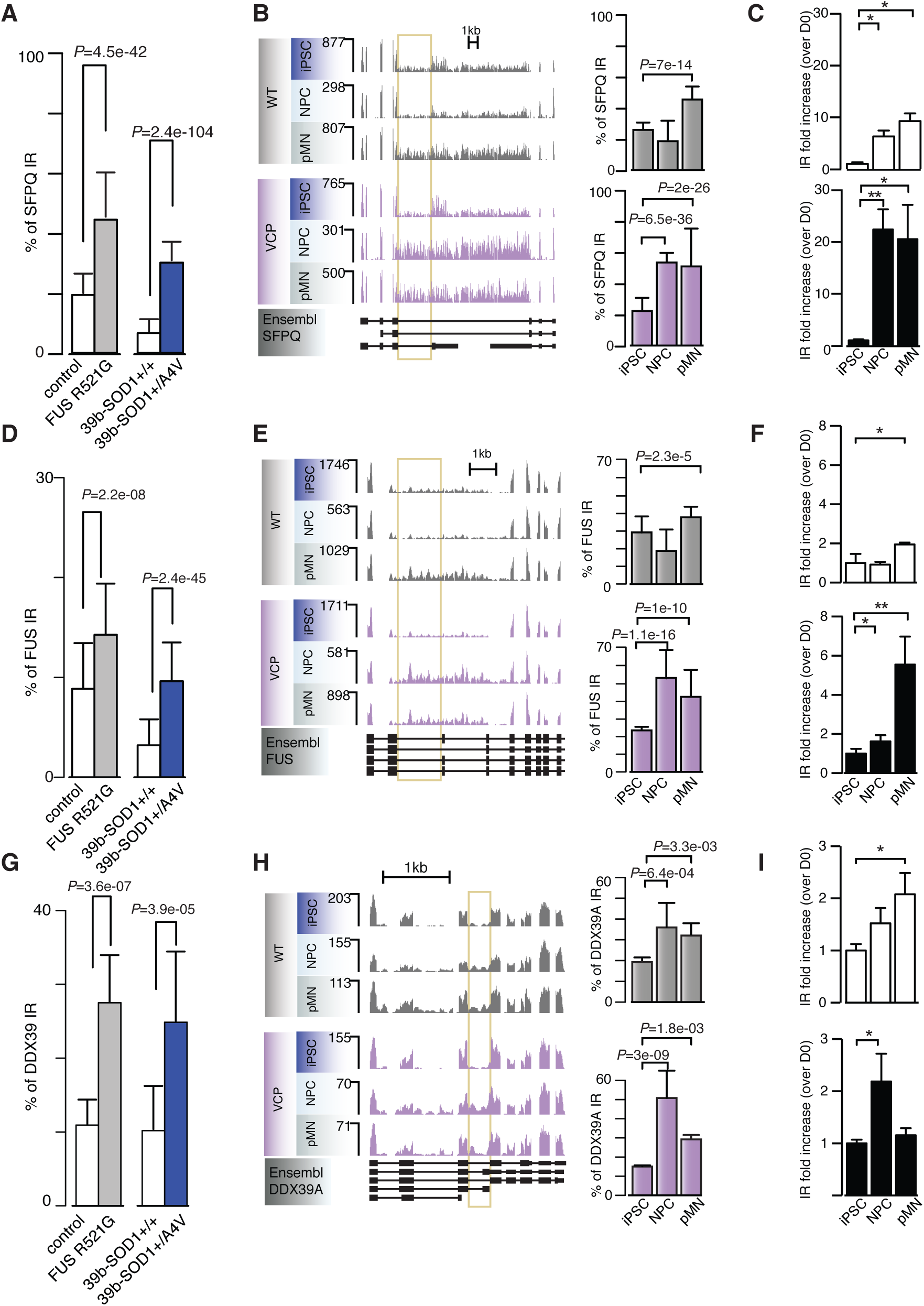
The most statistically significant IR event identified is in SFPQ, a splicing regulator. **(A)** Percentage intron retention in control compared with FUS*^mu^* or SOD1*^mu^* MNs for the gene SFPQ (mean ± SD; Fisher count test). **(B)** (*left*) Visualization of the RNA-seq read profiles of the intron-retaining gene SFPQ in control and VCP*^mu^* samples at iPSC, NPC and pMN stages. Intron of interest is indicated with yellow bar. (*right*) Bar graphs quantifying percentage intron retention across the entire time-course in controls and VCP samples (mean ± SD; Fisher count test). **(C)** Intron retention levels measured by qPCR. Levels of IR were measured using primer pair F2R2 (Figure S3) and were normalised over the gene’s expression levels (primer pair F1R1). To evaluate changes in the levels of IR over time within a group, IR levels at every timepoint are directly compared to those at the iPSC stage of the same group. N=3 control lines and N=4 VCP lines (mean+SD. * p<0.05, **p<0.01, one-way ANOVA with Dunnet correction for multiple comparisons). (**D-F**) Same as **A-C** for gene FUS. (**G-I**) Same as **A-C** for gene DDX39. See also Figure S3.

### Peaks of transient 3’ UTR remodelling and IR occur sequentially in early neurogenesis

Splicing and polyadenylation regulation are often interconnected. To examine whether 3’ UTR length varies throughout the distinct stages of MN differentiation and how the VCP mutation affect this process, we first extended the current catalogue of Ensembl 3’ UTR annotation using our RNA-seq data (as detailed in the methods). We then analysed their maximum expressed length over the course of differentiation. As expected, the 12,364 genes reliably expressed across all time-points exhibit longer 3’ UTR in mMNs compared to iPSC for both control and VCP*^mu^* samples. Notably, however, these exhibit shorter 3’ UTR in NPC compared to iPSC in control samples only (P-value < 0.01; Wilcoxon test; Figure 4A).

**Figure 4.**
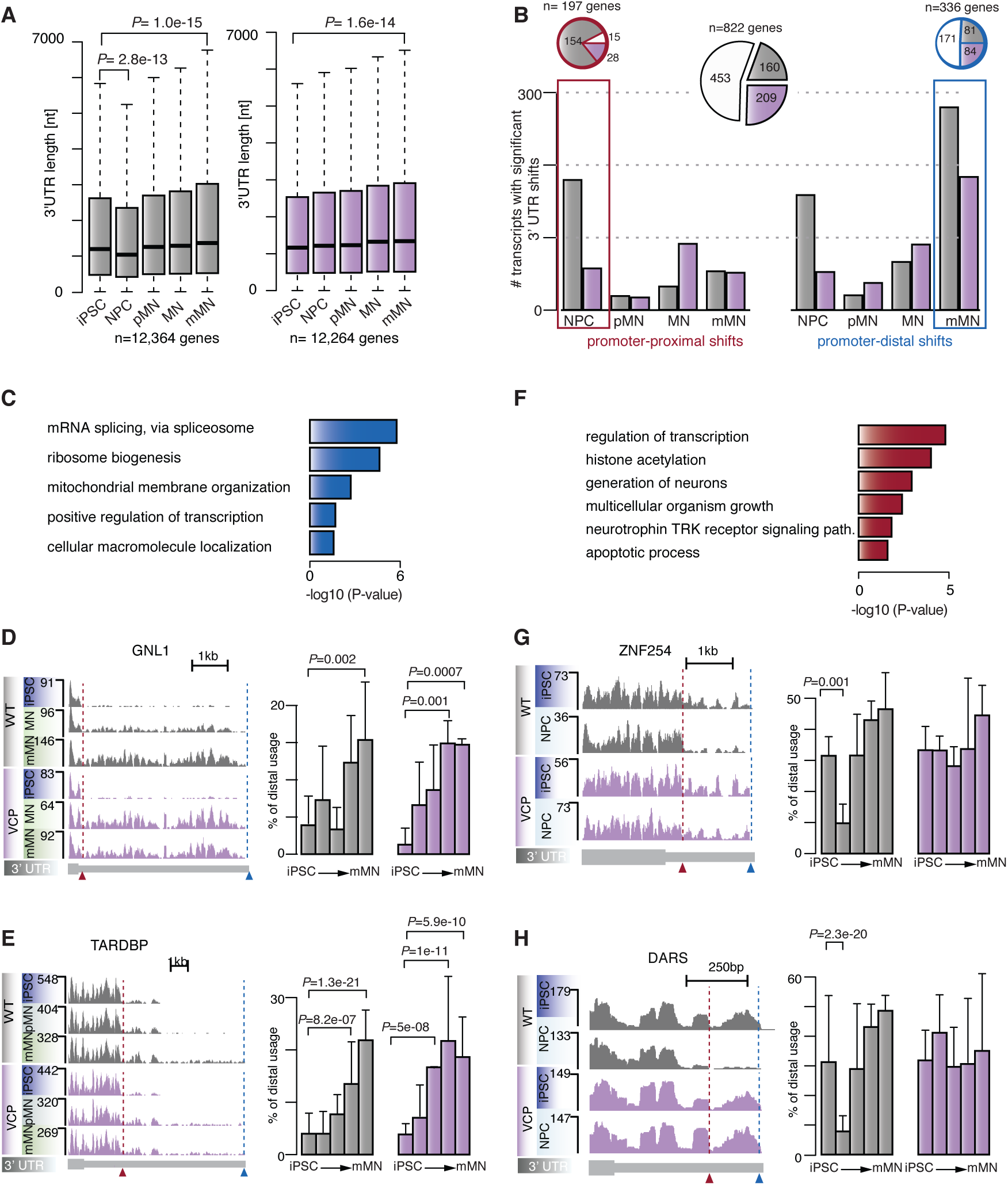
3’ UTR length variation during human motor neurogenesis. **(A)** Maximum 3’ UTR length of Ensembl transcript IDs expressed at distinct stages of MN differentiation in control (*left*) and VCP samples (*right*). Data shown as box plots in which the centre line is the median, limits are the interquartile range and whiskers are the minimum and maximum; *P*-value obtained with Wilcoxon test. **(B)** Number of 3’ UTR with statistically significant promoter-proximal (*left*) and promoter-distal (*right*) shifts at distinct stage of differentiation compared with iPSC in control (grey bars) and VCP samples (magenta bars). (*Inset*) Pie chart representing the proportions of genes exhibiting alternative 3’ UTR usage over time in control and VCP samples (white), control samples only (grey area) or VCP samples only (magenta area). **(C)** GO enrichment analysis of biological pathways associated with genes showing statistically significant distal shifts in poly(A) site usage in control mMN compared to control iPSCs. **(D** and **E)** (*left)* Genome browser views of the 3’ UTRs of genes GNL1 and TARDBP exhibiting statistically significant shifts towards increased promoter-distal poly(A) site usage in mMN compared with iPSCs. (*Right*) Bar plots showing the distal 3’ UTR usage relative to short 3’ UTR. *P-*value obtained from Fisher count test. **(F)** Same as **C** for genes showing statistically significant proximal shifts in poly(A) site usage in control NPCs compared with control iPSCs. **(G** and **H)** Same as **D** for the genes ZNF254 and DARS exhibiting statistically significant shifts towards increased promoter-proximal poly(A) site usage in NPC compared with iPSC. See also Figure S4.

To investigate further the dynamics of 3’ UTR processing, we analysed tandem poly(A) sites that were located in the same terminal exon. We used the number of reads mapped to the terminal 100 nt segment of each 3’ UTR as a proxy for the expression level of a 3’ UTR isoform to analyse the relative use of alternative poly(A) sites in different conditions. Overall we observed that 822 genes undergo significant 3’ UTR changes across the entire time-course of differentiation in either control and/or VCP*^mu^* cultures (Figure 4B). In 171 of these genes, we observed increased usage of distal poly(A) sites in both control and VCP*^mu^* mMNs (Figure 4B), consistent with a previous study (Ji et al., 2009).(Ji et al., 2009)W(Ji et al., 2009) also found that genes with statistically significant 3’ UTR lengthening in MNs were enriched in GO terms representing mRNA splicing (Figure 4C). About 50% of the genes exhibiting 3’ UTR lengthening from iPSC to mMNs were shared between control and VCP*^mu^* samples (Figure 4B) with <10 genes displaying significant proximal or distal 3’ UTR shifts when directly comparing control to VCP cultures at the at MN stage (Figure S4A). Although the ALS-causing VCP-mutation overall leads to similar 3’ UTR lengthening upon terminal differentiation as control samples, it is noteworthy that in several cases, VCP*^mu^* samples exhibited a significant promoter-distal shift at a comparatively earlier stage to control samples. These examples include GNL1 (Figure 4D) and TARDBP (Figure 4E), the later encoding Transactive-response DNA-binding Protein, 43 kDa (TDP-43) a splicing regulator, which becomes mislocalized and aggregated to form the pathological hallmark in > 95% of all ALS cases (Neumann et al., 2006), including VCP-related ALS (Johnson et al., 2010).

Surprisingly, 169 of the 822 genes exhibited statistically significant promoter-proximal shift specifically in control NPCs compared to iPSCs (Figure 4B). As 3’ UTR lengthening is expected upon embryonic development, the extent of 3’ UTR shortening at NPC stage is surprising. Various biological pathways such as transcription and neurotrophin regulation are represented among the transcripts exhibiting transient 3’ UTR shortening (Figure 4F). The majority of events are restricted to control samples, as only ∼20% of the above genes exhibiting 3’ UTR shortening at NPC stage only were found in VCP mutants. (Figures 4B, G and H). This is further confirmed when directly comparing VCP to control samples at the NPC stage where >200 promoter-distal shifts were detected (Figures S4A and S4B). It is interesting to note that the few 3’ UTR-shortening events at the VCP*^mu^* NPC stage include the gene HTT, which is required for neuronal development (Kerschbamer and Biagioli, 2015) (Figure S4C). These data demonstrate that extensive 3’ UTR length variation peaks prior to IR and characterises the transition from iPSCs to NPCs. Importantly, VCP samples do not exhibit any apparent similar process, which may either be explained by its absence or premature occurrence, which therefore escapes detection in our experimental paradigm.

### Transcriptional programs underlying human motor neurogenesis remain unperturbed by the VCP-mutation

Having identified major differences in post-transcriptional remodelling in ALS samples compared to control, we next sought to understand their functional consequences. Given the finding that VCP mutation deregulates the temporal dynamics of RNA processing during MN differentiation, it is surprising that ALS genetic background does not drive measurable transcriptional differences during motor neurogenesis **(Figure 1B).** Noting that hierarchical clustering is often dominated by a single transcriptional program, we applied singular value decomposition to the expression matrix (15,989 genes across 31 samples) to identify major orthogonal transcriptional programs and their associated biological processes underlying motor neurogenesis (Luisier et al., 2014). 69% of the variance in gene expression was captured by the first three components (Figure S5A) which we display in Figures 5A-C. Gene Ontology (GO) functional enrichment analysis of the groups of genes that were either positively or negatively associated with these components shows that cell proliferation and neurogenesis functions dominate component 1 (47% of gene expression variation). Components 2 and 3 (15% and 7% of gene expression variation respectively), are dominated by more specific biological pathways such as regionalization, patterning and axonogenesis (Figures 5D-F). These results highlight the orthogonal cellular components that operate in parallel: progressive neurogenesis (component 1), transient neural induction and patterning (component 2) and terminal differentiation with some contribution to induction (component 3). Importantly, control and VCP-mutant samples behave similarly in these first three components. A multivariate linear analysis confirmed that time rather than VCP mutation is the most explanatory variable in components 1, 2 and 3 (Figure S5B). In summary, we find that transcriptional programs reflect the developmental trajectory of MNs as specific pathways are activated in a developmental stage-dependent manner. However these developmental programs are not disrupted by the VCP mutation despite post-transcriptional defects, suggesting an underlying robustness of these transcriptional programs and/or compensatory mechanisms.

**Figure 5.**
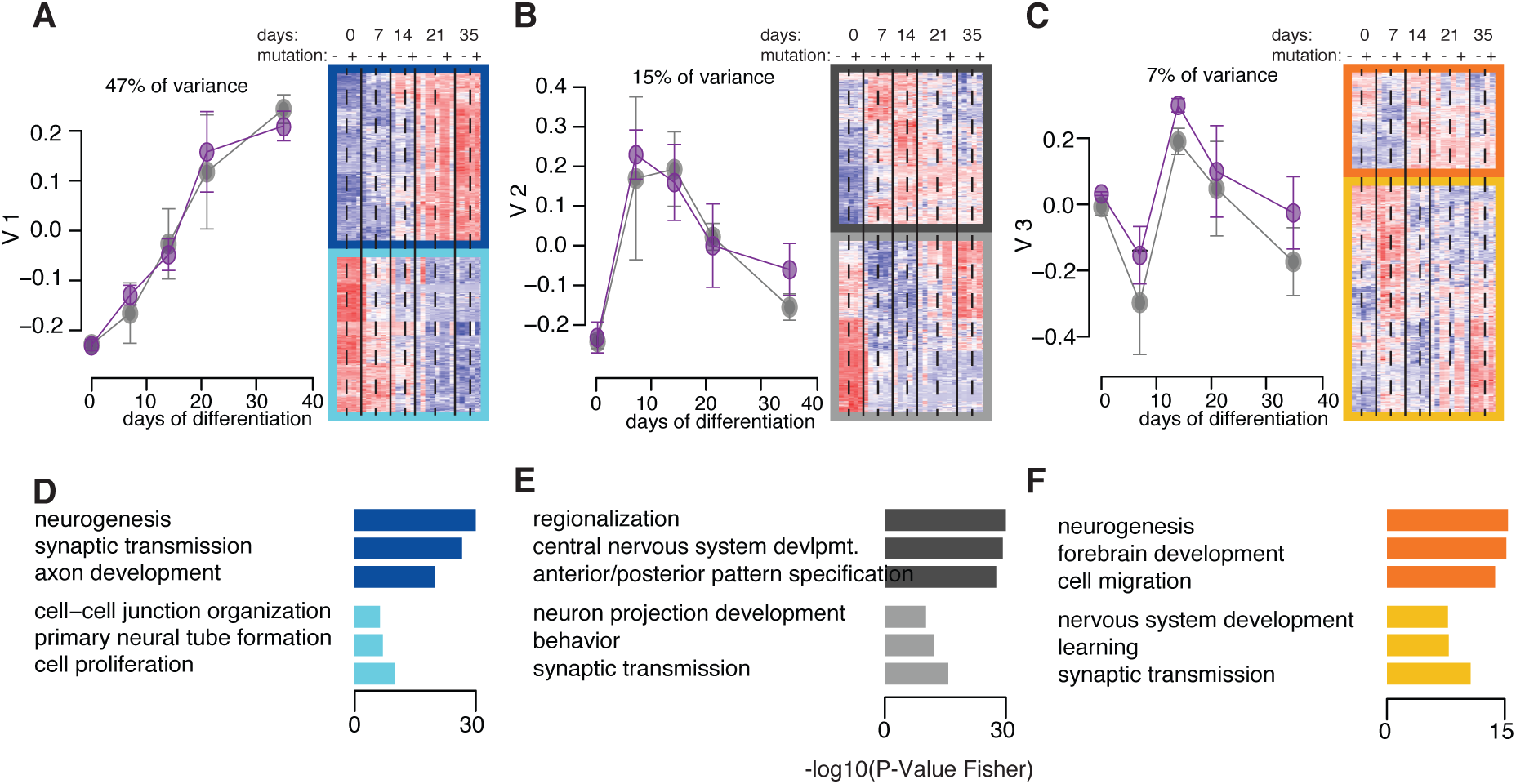
The integrity of transcriptional programs underlying human motor neurogenesis is preserved despite the VCP-mutation. **(A-C)** Singular value decomposition analysis of the expression of 15,989 genes in n = 31 samples. (*left*) Expression profiles of the first three singular vectors 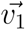 through 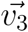, capturing respectively 47%, 15% and 7% of the variance in gene expression. Grey and magenta points indicate expression for the control samples and for the VCP samples respectively. (*right*) Heatmap of the standardized expression of the genes contributing and correlating positively (top 3 darker rectangles) and negatively (bottom 3 lighter rectangles) with the first three right singular vectors. **(D-F)** For each of the first three components a selection of biological pathways (Gene ontology) that are enriched among the genes contributing and correlating positively (top 3 rectangles) or negatively (bottom 3 rectangles) with the first three right singular vector is plotted below the expression profiles. The size of each bar corresponds to the -log10(P-value) of enrichment. See also Figure S4.

## DISCUSSION

The objectives of this research study were twofold: to resolve the alternative RNA processing events underlying distinct stages of MN lineage restriction and systematically examine the influence of ALS-causing VCP mutation on this process. In order to achieve this, we integrated the directed differentiation of patient-specific iPSCs into spinal MNs with RNA sequencing and comprehensive bioinformatic examination. We show that a robust splicing program underlies MN development as summarised in Figure 6. In resolving the precise nature of sequential post-transcriptional programmes underlying distinct stages of MN differentiation, we show that the timing of these carefully choreographed molecular events is perturbed by the ALS-causing VCP mutation. In analysing additional ALS-related MNs samples, we further show that ALS gene backgrounds such as SOD1 and FUS affect the transcript structure of key RNA splicing regulators in a similar manner to those targeted by the accelerated splicing program in VCP samples. Notably these include SFPQ, a key regulator of axonal mRNA localization and axonal development which dysregulated transcript structure may play a key role in axonal degeneration during ALS development. Indeed SFPQ is emerging as a key regulator of neurodegeneration more widely (Cosker et al., 2016; Ishigaki et al., 2017; Ke et al., 2012; Thomas-Jinu et al., 2017). Finally we show that transcriptional programs underlying human motor neurogenesis remain unperturbed by the VCP-mutation in spite of post-transcriptional defects.

**Figure 6.**
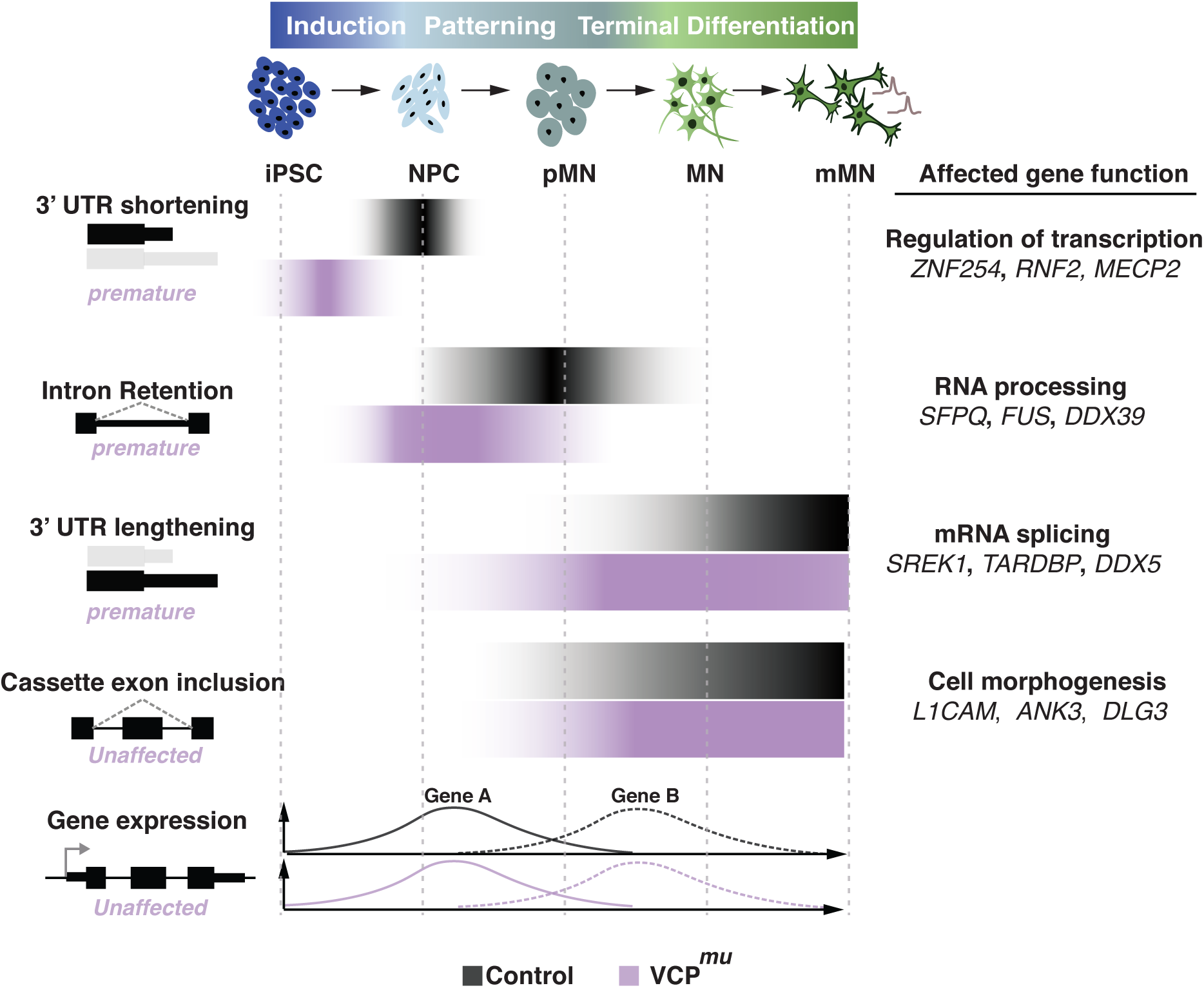
Proposed model. Cartoon summarising the kinetics of post-transcriptional events underlying human motor neurogenesis from control and VCP*^mu^* iPSCs. Our results suggest a model where the predominant RNA processing events at an early stage of neural differentiation are IR and 3’ UTR remodelling. These events start prematurely in VCP*^mu^* samples but do not affect major transcriptional programs underlying human MN differentiation. Conversely, cassette exon inclusion exhibits similar kinetics in control and VCP*^mu^* samples.

### The temporal dynamics of post-transcriptional events underlying human motor neuron development

Dynamic AS changes have been previously reported in rodent forebrain between different developmental stages (Dillman et al., 2013; Yan et al., 2015). Here we have identified previously unrecognised modifications in the transcript structure of genes regulating RNA processing and splicing at early stages of human MN lineage restriction (Figure 6). We show that a peak of 3’ UTR remodelling, including 3’ UTR shortening, precedes IR. Specifically we show that significant 3’ UTR length variation affects transcript structure as cells exit pluripotency during neural induction. This precedes the peak of intron-retaining transcript accumulation that occurs upon neural patterning to the ventral spinal cord. Following this, progressive 3’ UTR lengthening, cassette exon inclusion and intron splicing then characterise the remaining phases of terminal differentiation to MNs as previously shown (Yap et al., 2012; Zhang et al., 2016).

### Transient 3’ UTR remodelling characterises neural induction

3’ UTRs play key roles in post-transcriptional gene expression regulation and mRNAs expressed in brain tissues generally have longer 3’ UTRs compared to other tissues (Miura et al., 2013; Wang and Yi, 2014; Zhang et al., 2005). Indeed it is also established that progressive lengthening of mRNA 3’ UTRs is observed during mouse embryonic development (Ji et al., 2009). Here we show extensive 3’ UTR length variation during neural induction from human iPSCs. Unexpectedly, we found significant 3’ UTR transient shortening that precedes the - previously reported - progressive 3’ UTR lengthening towards terminal differentiation. The molecular function of such 3’ UTR shortening remains unclear, although other studies report a potential role of 3’ UTR in paraspeckle formation and the exit of pluripotency (Bond and Fox, 2009; Chen and Carmichael, 2009; Naganuma et al., 2012). The highly transient nature of this regulatory process may explain why it has evaded experimental detection to date. Our time-resolved experimental paradigm has permitted the identification of sequential transcriptional programmes underlying human motor neurogenesis.

### Transient IR characterises an early stage of MN lineage restriction

IR is emerging as novel post-transcriptional mechanism regulating gene expression. Neural and immune cell types have higher proportions of retained introns compared to other tissues (Braunschweig et al., 2014). Previous studies have demonstrated progressive IR during terminal differentiation of mouse neurons and the relevance of intron-retaining transcripts in regulating the expression of functionally linked neuron-specific genes in mouse models (Braunschweig et al., 2014; Yap et al., 2012). Here we show evidence for a transient ‘wave’ of intron-retaining transcripts at pMN stage that targets a network of splicing regulators. Recognizing the interaction between IR events and nuclear paraspeckles, this raises the possibility that either these transcripts contribute to the stabilisation of the paraspeckles or that they are dynamically sequestered in such nuclear structures in order to transiently inhibit their function. Further experiments are required to determine the stability and localisation of such transcripts.

### Overall relevance of aberrant IR to ALS

We show that the timing of the post-transcriptional program is perturbed in samples with ALS-causing VCP mutation, with a premature IR and to some extent 3’ UTR length variation. Interestingly IR was the most prominent change in transcript structure at early stages of lineage restriction and also exhibited striking premature initiation in VCP mutants. Additionally key RNA splicing regulators that were targeted by the accelerated splicing program in VCP*^mu^* samples also exhibited widespread IR in MNs harbouring mutations in other ALS-causing genes, including SOD1 and FUS.

While the accelerated splicing program in VCP-mutant samples might be linked to neuronal degeneration, the lack of transcriptional changes indicates that the dysfunctions in IR and 3’ end processing do not affect neuronal development in a major way. Thus, the post-transcriptional changes appear to be compensated during the process of neuronal differentiation. This raises the important possibility that early mutation-dependent changes in post-transcriptional remodelling may induce a state of compensated dysfunction, which could culminate in a disease phenotype later once the compensatory mechanisms are diminished. We have previously shown that the phenotype of VCP mutation can be detected only once MNs have begun extending axons (Hall et al., 2017), indicating that the process of differentiation may sensitise the MNs to the pre-existing post-transcriptional defects. Therefore it is reasonable to think that accelerated IR represents subtle manifestations of a dysregulated pathway of the developing neurons, which eventually contributes to the increased vulnerability of the differentiated MNs.

Notably the most significant intron-retaining transcript in ALS samples was SFPQ, which encodes a protein that plays key roles in salient ALS-related pathways including RNA transcription, splicing, 3’ end processing and axon viability (Arnold et al., 2013; Clark et al., 2016; Cosker et al., 2016; Danckwardt et al., 2007; Dong et al., 2007; Hall-Pogar et al., 2007; Hirose et al., 2014; Masuda et al., 2016; Patton et al., 1993). SFPQ IR at early stages of MN development is required to ensure the integrity of downstream splicing events, as suggested by IR in control motor neurogenesis. SFPQ IR may however be detrimental at the terminally differentiated MN stage where it plays key role in regulating mRNA axonal transport and axon viability (Cosker et al., 2016; Thomas-Jinu et al., 2017). Additionally SFPQ together with other proteins is required for nuclear paraspeckle integrity. It is therefore reasonable to speculate that SFPQ IR in MN might contribute to paraspeckle formation that has been previously observed at an early stage of the ALS pathogenesis (Lourenco et al., 2015; Nishimoto et al., 2013; Sasaki et al., 2009). SFPQ plays diverse roles at different stages of MN differentiation and therefore IR may have context-dependent functional consequences.

Collectively, these findings are consistent with a model whereby ALS-causing mutations lead to aberrant IR and 3’ end processing during motor neurogenesis, which may be a primary molecular event that ultimately contributes to the disruption of motor neuron integrity. Our findings also indicate that these early perturbations in post-transcriptional processing are well compensated, but this state of compensated dysfunction increases the vulnerability of differentiated MNs towards initiating disease onset and progression.

## AUTHOR CONTRIBUTIONS

Conceptualization, R.L., J.U., N.M.L., R.P; Formal Analysis, R.L.; Investigation, R.L., G.E.T., C.E.H.; Writing – Original Draft, R.L., R.P.; Writing – Review & Editing, R.L., J.U., N.M.L., R.P.; Resources, R.L., J.U., N.M.L., R.P.; Visualization, R.L.; Funding Acquisition, R.L., J.U., N.M.L., R.P.; Supervision, J.U., N.M.L., R.P.

## ACKNOWLEDGMENTS

We are thank Miha Modic for sharing with us RNA-seq data from hESC. We thank Martina Hallegger and Roberto Simone for their input regarding primer designs. The authors wish to thank the patients for fibroblast donation. This work was supported by the Francis Crick Institute which receives its core funding from Cancer Research UK (FC010110), the UK Medical Research Council (FC001002), and the Wellcome Trust (FC001002), the Wellcome Trust funding to R.P. [101149/Z/13/A], to N.L. and J.U. [103760/Z/14/Z], Takeda Cambridge and Cerevance (R.P), Grand Challenges (C.E.H.), a Marie Curie Post-doctoral Research Fellowship 657749-NeuroUTR (R.L.), and an Advanced Postdoc Mobility fellowship from the Swiss National Science Foundation P300PA_174461 (R.L.).

## Supplementary Figure 1. Related to Figure 1

**Supplementary Figure 1. Related to Figure 1.**
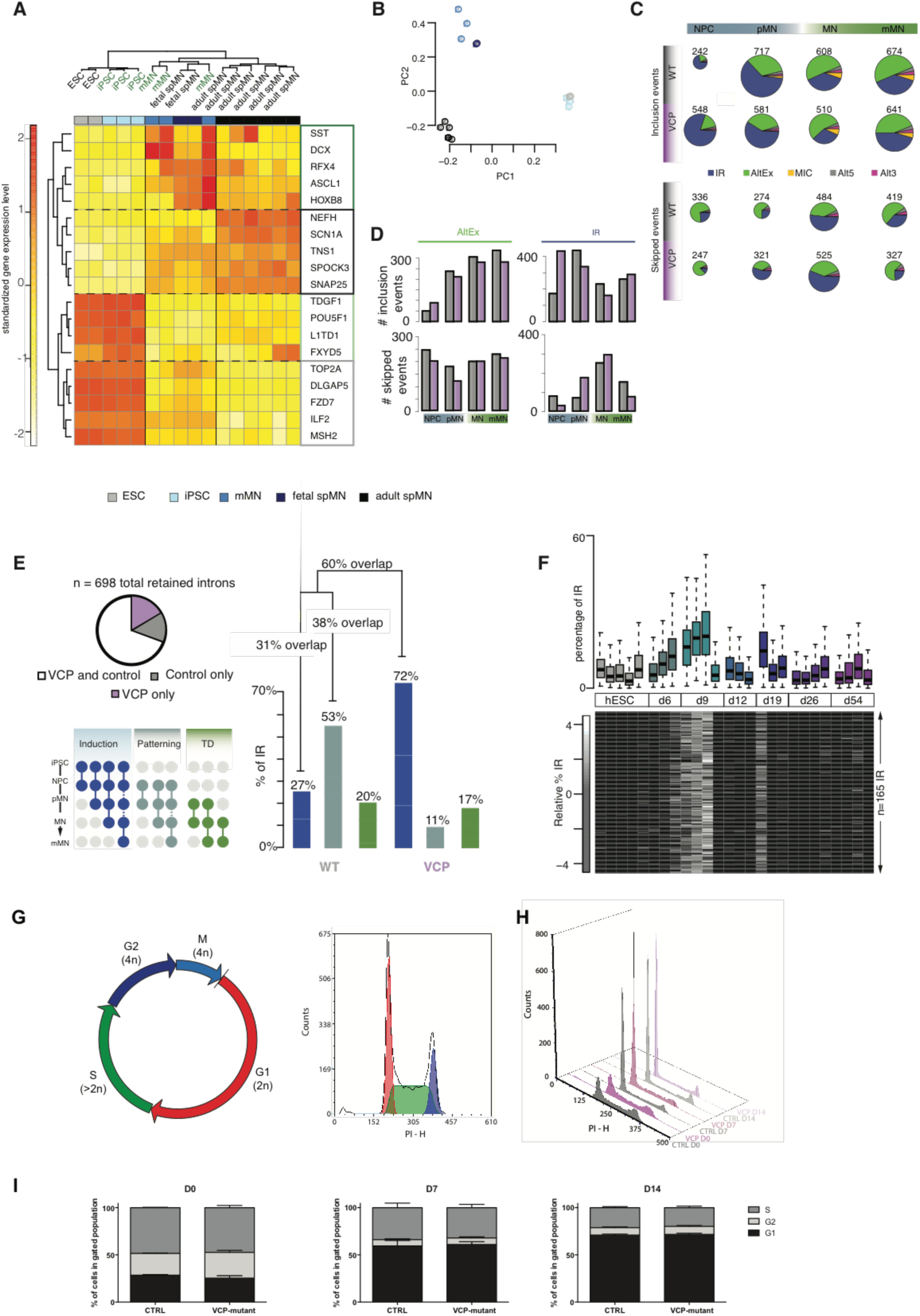
**(A)** Heatmap of the standardized expression of 19 key gene markers of spinal MN (spMN) maturation and embryonic development identified in a previous study (Ho et al., 2016). Samples in green come from the present study. Two ESC RNA-seq data were obtained from collaborators (unpublished data). Two fetal spMN (NIH Roadmap Epigenomics Mapping Consortium) and two adult spMN (laser-captured spMN; (Nichterwitz et al., 2016)) samples were downloaded from Array Express and GEO. **(B)** PCA performed on normalized gene expression values of key gene markers. Samples are plotted by their coordinates along PC1 and PC2. Colors of data points indicate similar sample types. ESC (grey), iPSC (light blue), mMN (blue), fetal spMN (navy) and adult spMN (black). **(C)** Pie charts representing distributions of regulated included (*upper*) and skipped (*lower*) splicing events in healthy control and VCP-mutant samples at distinct stages of motor neurogenesis compared to iPSCs or previous time-point. Intron retention (IR); alternative exon (AltEx); micro-exons (MIC); alternative 5’ and 3’ utr (Alt5 and Alt3). **(D)** Bar graphs representing the number of included (*upper*) and skipped (*lower*) exonic and intronic splicing events respectively in healthy controls (grey bars) and VCP samples (magenta bars) at specific time points during motor neuron differentiation. **(E)** (*upper left*) Pie chart representing percentages of retained introns across all time points that are either shared between VCP and control samples (white) or specific to either control (grey) or VCP samples (magenta). (*lower left*) Schematic showing colour codes for different categories of IR grouped according to stage of initiation and persistence. Coloured circles indicate sample status and line between circles indicate event statistically significant between sample and either prior sample from prior status or iPSC. (*right*) Bar graphs representing the percentage of intronic retention events upon induction, patterning or terminal differentiation in healthy control or VCP samples. The heights of the different coloured stacked bars represents the relative number of IR events that last until specific stages of lineage restriction as shown on lower left part of the panel. Percentage of overlap between events in control and VCP events is shown above the stacked bars. **(F)** Percentage of retention for 167 manually curated introns in a study of in vitro neural differentiation of hESCs for 54 days (Yao et al., 2017). **(G)** Diagram showing cell cycle and its analysis. DNA content of each cell is quantified by measuring the intensity of propidium iodide (PI) staining by flow cytometry. Cells in G2 (tetraploid, 4n, blue) have a two times higher PI intensity than cells in G1 (diploid, 2n, red). Cells in the S phase are replicating their genetic material and have intermediate PI intensity (>2n, green). A representative flow cytometry histogram with automated interpolation of each phase is shown on the right. **(H)** Representative cell cycle histograms for both control and VCP*^mu^* lines at each time point. **(I)** At each time point the proportion of cells in either G1 (black), S (grey) or G2/M (light grey) were automatically calculated. Data is expressed as mean+SD, N=4 control lines vs N=4 VCP-mutant lines from two independent experiments (i.e. two independent neural inductions).

**Supplementary Figure 2. Related to Figure 2.**
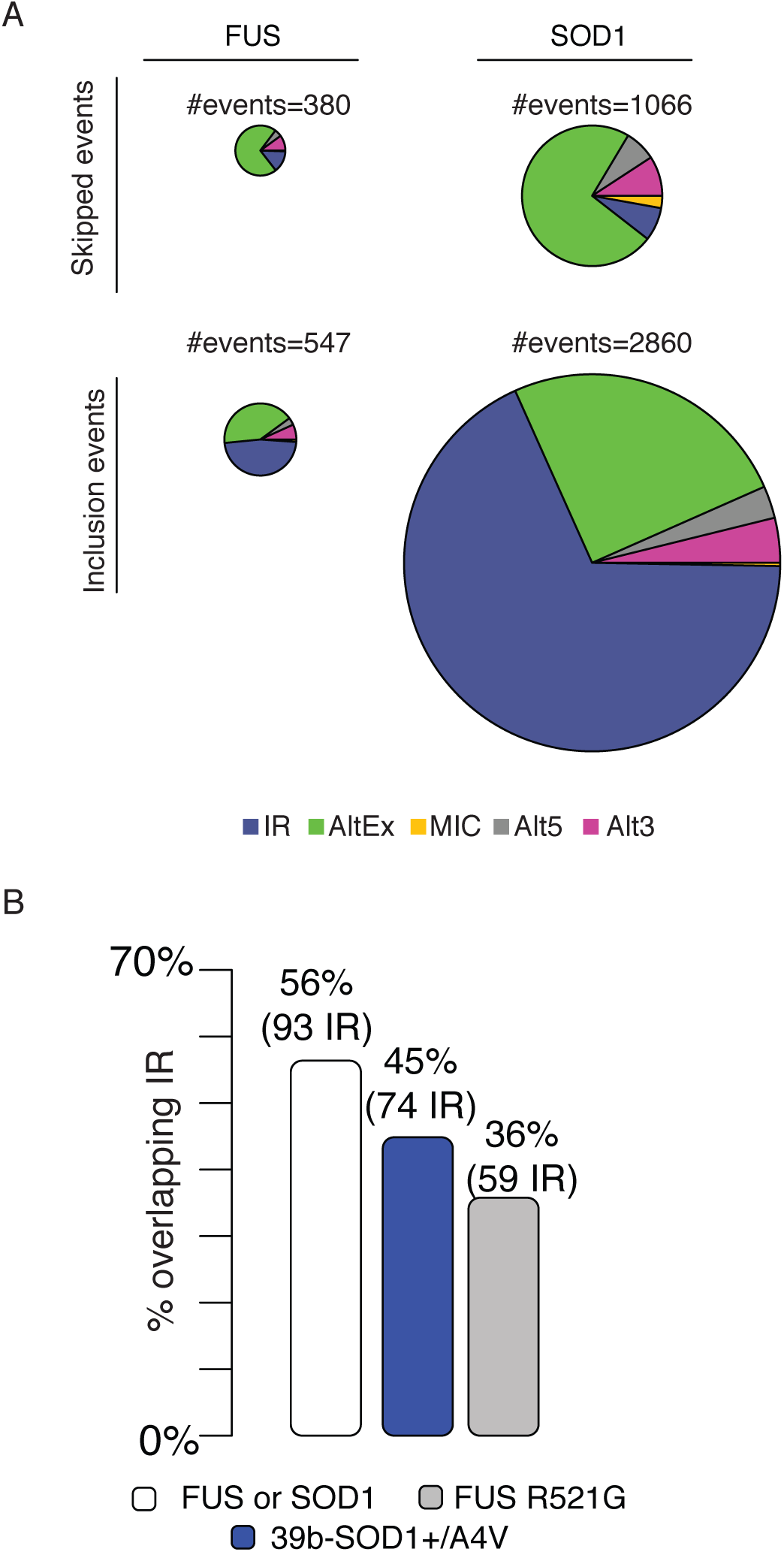
(A) Pie charts representing distributions of regulated splicing events in SOD1 and FUS mutant MNs compared to controls. Intron retention (IR); alternative exon (AltEx); micro-exons (MIC); alternative 5’ and 3’ utr (Alt5 and Alt3). (B) Bar graphs depicting the percentage of retained introns upon MN differentiation identified in this study (167 events in 143 genes) that also exhibit significant retention in MN samples harbouring either SOD1 or FUS gene mutations (white bar), in SOD1-mutant MNs (blue bar) or FUS-mutant MNs (grey bar) (Kapeli et al., 2016; Kiskinis et al., 2014b).

**Supplementary Figure 3. Related to Figure 3.**
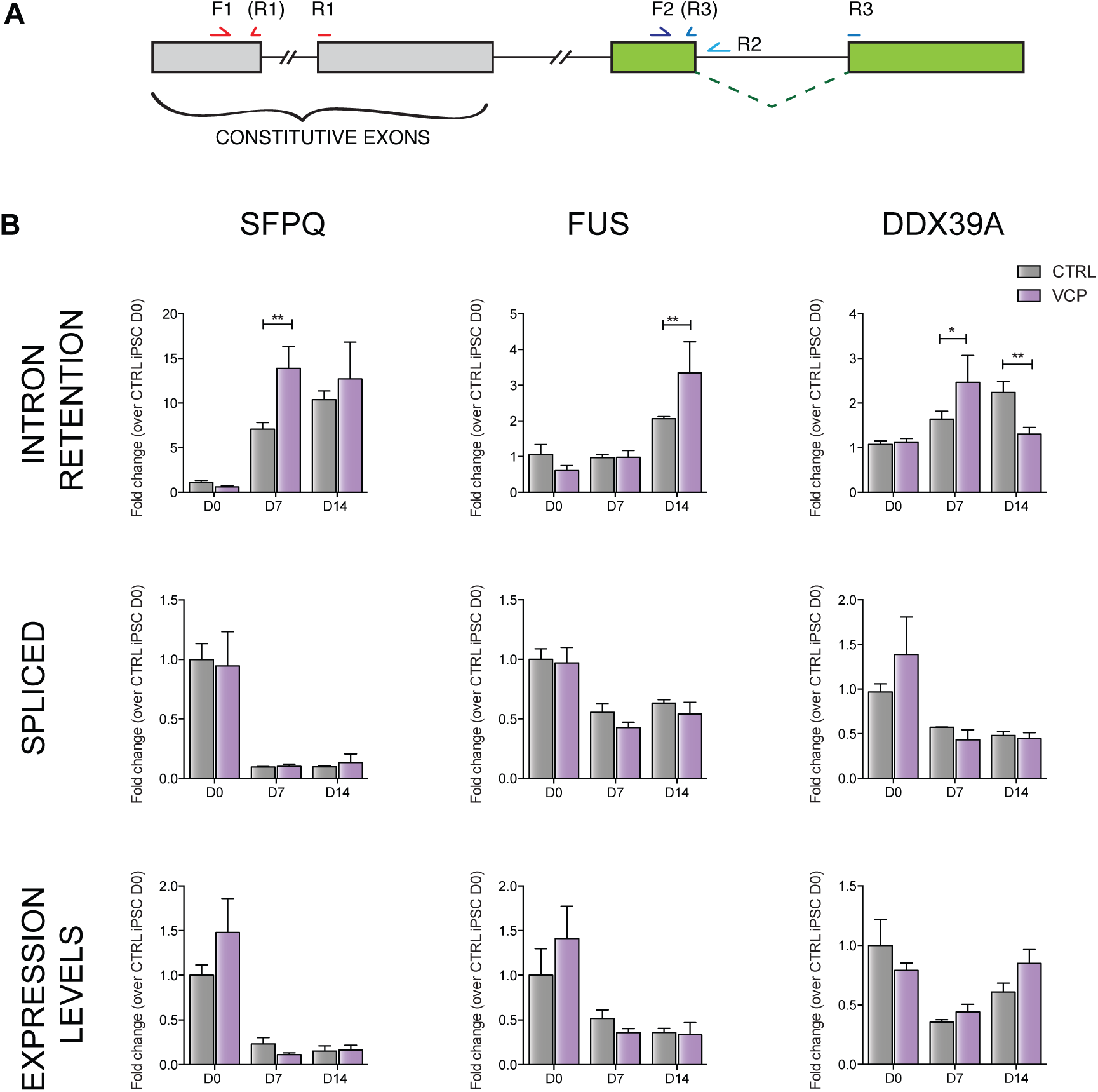
**(A)** Experimental design for qPCR validation of intron retention. For each transcript to be analysed, three primer pairs were used. Primer pair F1 R1 (intron spanning, across exon-exon junction) were used to to analyse gene expression levels. Primer pair F2 R2 (one primer on an exon flanking the intron to be analysed, the other on the intron) was used to assess the levels of intron retention. Primer pair F3 R2 (both primers on the exons flanking the intron of interest, if possible designed across the exon-exon junction) was used to measure levels of the spliced transcript. **(B)** Intron retention levels measured by qPCR. Levels of IR were measured using primer pair F2R2 and were normalised over the gene’s expression levels (primer pair F1R1). Gene expression levels were normalised using three housekeeping genes (GAPDH, POLR2B and UBE2D3). To compare levels of IR at any given time point across groups, IR levels were normalised over IR levels in control cells at D0. N=3 control lines and N=4 VCP lines. Data is expressed as mean+SD. *p<0.05, **p<0.01, two-way ANOVA with Sidak’s correction for multiple comparisons.

**Supplementary Figure 4. Related to Figure 4.**
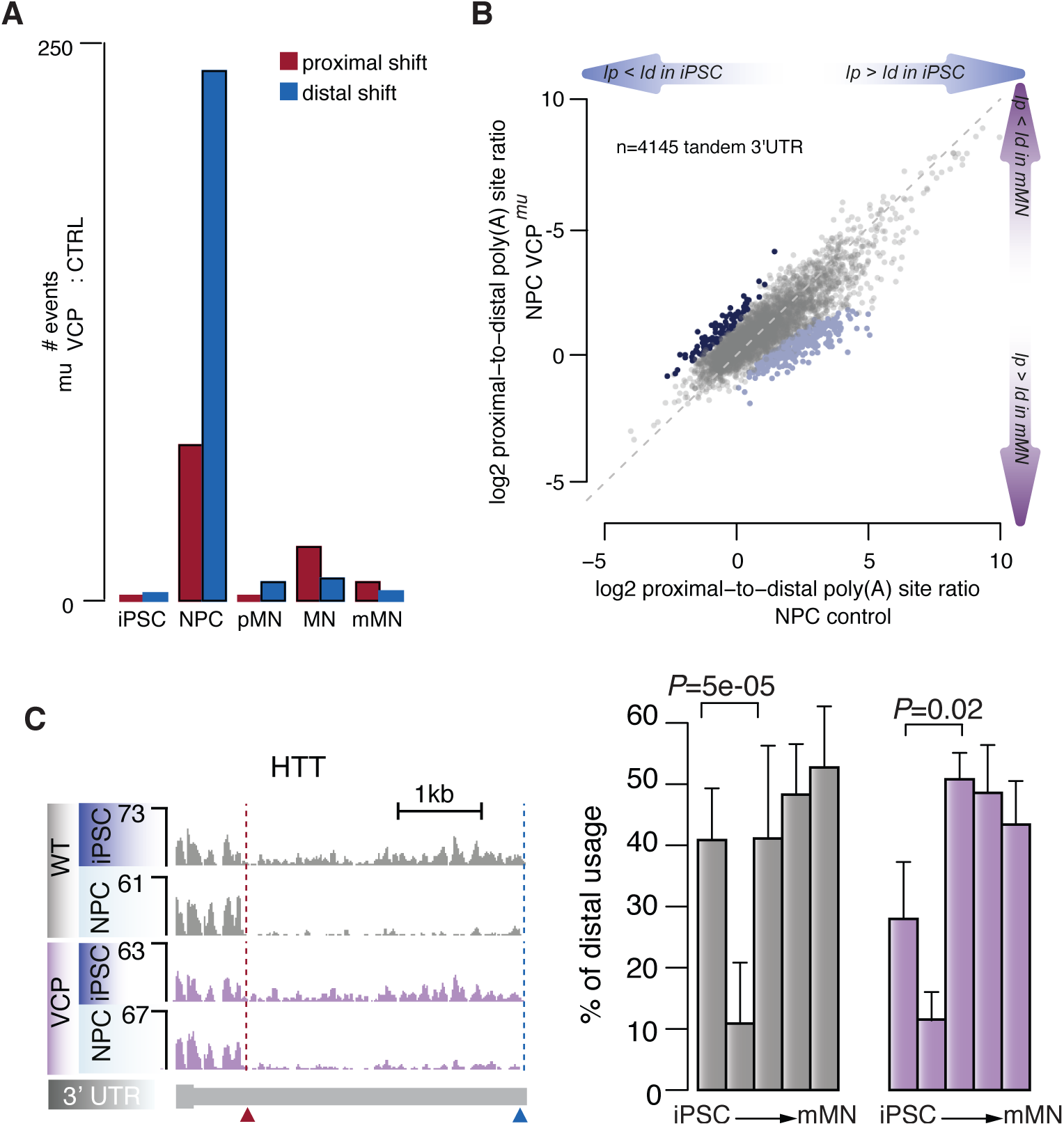
**(A)** Bar plot showing number of 3’ UTR with significant promoter-proximal (*red*) and promoter-distal (*blue*) shifts at each stage of differentiation in VCP samples compared to control samples. **(B)** Scatter plot of the relative use of promoter-proximal and promoter-distal poly(A) sites in control and VCP samples at NPC stage. FDR<0.01 between VCP and control NPC samples (Fisher exact test). Dark blue = promoter-proximal shifts in VCP compared to control. Light blue = promoter-distal shifts in VCP compared to control. **(C)** Genome browser view of the 3’ UTR of the genes HTT exhibiting significant shift towards increased promoter-proximal poly(A) site usage in NPC compared to iPSC. (*Right*) Bar plots showing the distal 3’ UTR usage relative to short 3’ UTR. *P-*value obtained from Fisher count test.

**Supplementary Figure 5. Related to Figure 5.**
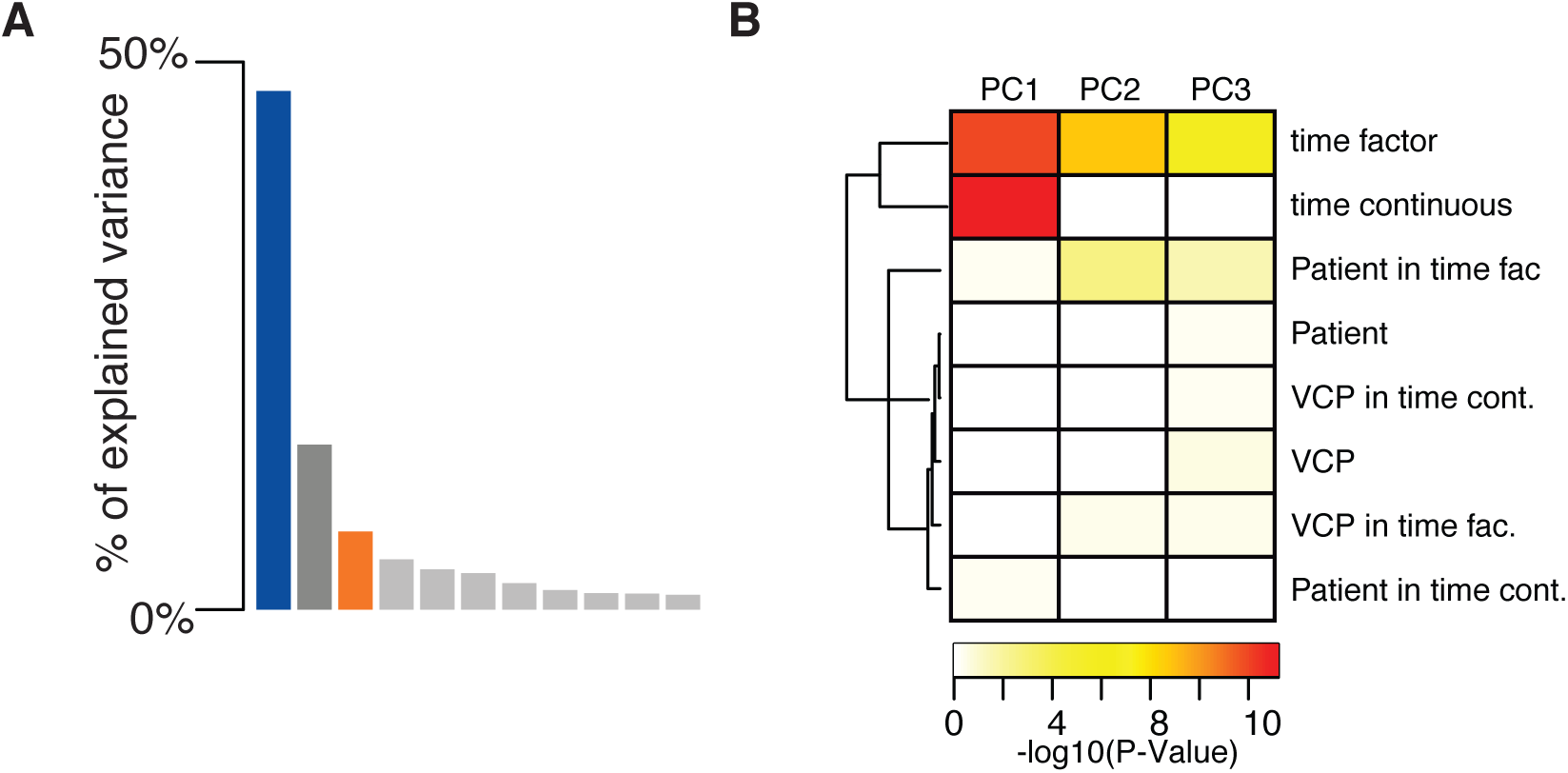
(A) Fraction of variance in gene expression captured by the corresponding principal components. (B) Heatmap of the significance of association between each variable in this study and the left singular vectors of the components 1, 2 and 3. Significance was obtained from P-value of the multivariate linear model.

**Table S1:** iPSC lines utilized in this study.

| iPSC line | Mutation present | Age of Donor | Age at disease onset | Sex of Donor |
| --- | --- | --- | --- | --- |
| CTRL 1 | None | 78 | N/A | Male |
| CTRL 2 | None | 64 | N/A | Male |
| CTRL 3 | None | (unknown) | N/A | Female |
| MUT 1 | R155C | 43 | 40 | Female |
| MUT 2 | R155C | 43 | 40 | Female |
| MUT 3 | R191Q | 42 | 36 | Male |
| MUT 4 | R191Q | 42 | 36 | Male |

**Table S2:** Metadata table of the RNA-seq samples generated in this study.

| FASTQ name | Sample Name | Source | Organism | Patient number | iPSC cell line | Technical replicate* | Mutation present | Days of differentiation | Age of Donor | Age at disease | Sex of Donor | GSE | Source |
| --- | --- | --- | --- | --- | --- | --- | --- | --- | --- | --- | --- | --- | --- |
| 1.fastq.gz | MUT 1 iPSC | iPSC cell | homo sapiens | Patient 1 | MUT 1 | 1 | VCP: R155C | 0 | 43 | 40 | Female | GSE98290 | this paper |
| 2.fastq.gz | MUT 2 iPSC | iPSC cell | homo sapiens | Patient 1 | MUT 2 | 1 | VCP: R155C | 0 | 43 | 40 | Female | GSE98290 | this paper |
| 3.fastq.gz | MUT 3 iPSC | iPSC cell | homo sapiens | Patient 2 | MUT 3 | 1 | VCP:R191Q | 0 | 42 | 36 | Male | GSE98290 | this paper |
| 4.fastq.gz | MUT 1 d7 | Neural precursor cells | homo sapiens | Patient 1 | MUT 1 | 1 | VCP: R155C | 7 | 43 | 40 | Female | GSE98290 | this paper |
| 5.fastq.gz | MUT 2 d7 | Neural precursor cells | homo sapiens | Patient 1 | MUT 2 | 1 | VCP: R155C | 7 | 43 | 40 | Female | GSE98290 | this paper |
| 6.fastq.gz | MUT 3 d7 | Neural precursor cells | homo sapiens | Patient 2 | MUT 3 | 1 | VCP:R191Q | 7 | 42 | 36 | Male | GSE98290 | this paper |
| 7.fastq.gz | MUT 1 d14 | pMN progenitor cells | homo sapiens | Patient 1 | MUT 1 | 1 | VCP: R155C | 14 | 43 | 40 | Female | GSE98290 | this paper |
| 8.fastq.gz | MUT 2 d14 | pMN progenitor cells | homo sapiens | Patient 1 | MUT 2 | 1 | VCP: R155C | 14 | 43 | 40 | Female | GSE98290 | this paper |
| 9.fastq.gz | MUT 1 d21 | electrically immature motor neurons | homo sapiens | Patient 1 | MUT 1 | 1 | VCP: R155C | 21 | 43 | 40 | Female | GSE98290 | this paper |
| 10.fastq.gz | MUT 2 d21 | electrically immature motor neurons | homo sapiens | Patient 1 | MUT 2 | 1 | VCP: R155C | 21 | 43 | 40 | Female | GSE98290 | this paper |
| 11.fastq.gz | MUT 2 d21 | electrically immature motor neurons | homo sapiens | Patient 1 | MUT 2 | 2 | VCP: R155C | 21 | 43 | 40 | Female | GSE98290 | this paper |
| 12.fastq.gz | MUT 3 d21 | electrically immature motor neurons | homo sapiens | Patient 2 | MUT 3 | 1 | VCP:R191Q | 21 | 42 | 36 | Male | GSE98290 | this paper |
| 13.fastq.gz | MUT 1 d35 | electrically mature motor neurons | homo sapiens | Patient 1 | MUT 1 | 1 | VCP: R155C | 35 | 43 | 40 | Female | GSE98290 | this paper |
| 14.fastq.gz | MUT 1 d35 | electrically mature motor neurons | homo sapiens | Patient 1 | MUT 1 | 1 | VCP: R155C | 35 | 43 | 40 | Female | GSE98290 | this paper |
| 15.fastq.gz | MUT 3 d35 | electrically mature motor neurons | homo sapiens | Patient 2 | MUT 3 | 1 | VCP:R191Q | 35 | 42 | 36 | Male | GSE98290 | this paper |
| 16.fastq.gz | CTRL 2 iPSC | iPSC cell | homo sapiens | Patient 3 | CTRL 2 | 1 | N/A | 0 | 78 | N/A | Male | GSE98290 | this paper |
| 17.fastq.gz | CTRL 1 iPSC | iPSC cell | homo sapiens | Patient 4 | CTRL 1 | 1 | N/A | 0 | 64 | N/A | Male | GSE98290 | this paper |
| 18.fastq.gz | CTRL 2 iPSC | iPSC cell | homo sapiens | Patient 3 | CTRL 2 | 2 | N/A | 0 | 78 | N/A | Male | GSE98290 | this paper |
| 19.fastq.gz | CTRL 2 d7 | Neural precursor cells | homo sapiens | Patient 3 | CTRL 2 | 1 | N/A | 7 | 78 | N/A | Male | GSE98290 | this paper |
| 20.fastq.gz | CTRL 1 d7 | Neural precursor cells | homo sapiens | Patient 4 | CTRL 1 | 1 | N/A | 7 | 64 | N/A | Male | GSE98290 | this paper |
| 21.fastq.gz | CTRL 2 d7 | Neural precursor cells | homo sapiens | Patient 3 | CTRL 2 | 2 | N/A | 7 | 78 | N/A | Male | GSE98290 | this paper |
| 22.fastq.gz | CTRL 2 d14 | pMN progenitor cells | homo sapiens | Patient 3 | CTRL 2 | 1 | N/A | 14 | 78 | N/A | Male | GSE98290 | this paper |
| 23.fastq.gz | CTRL 1 d14 | pMN progenitor cells | homo sapiens | Patient 4 | CTRL 1 | 1 | N/A | 14 | 64 | N/A | Male | GSE98290 | this paper |
| 24.fastq.gz | CTRL 2 d14 | pMN progenitor cells | homo sapiens | Patient 3 | CTRL 2 | 2 | N/A | 14 | 78 | N/A | Male | GSE98290 | this paper |
| 25.fastq.gz | CTRL 2 d21 | electrically immature motor neurons | homo sapiens | Patient 3 | CTRL 2 | 1 | N/A | 21 | 78 | N/A | Male | GSE98290 | this paper |
| 26.fastq.gz | CTRL 1 d21 | electrically immature motor neurons | homo sapiens | Patient 4 | CTRL 1 | 1 | N/A | 21 | 64 | N/A | Male | GSE98290 | this paper |
| 27.fastq.gz | CTRL 1 d21 | electrically immature motor neurons | homo sapiens | Patient 4 | CTRL 1 | 2 | N/A | 21 | 64 | N/A | Male | GSE98290 | this paper |
| 28.fastq.gz | CTRL 2 d21 | electrically immature motor neurons | homo sapiens | Patient 3 | CTRL 2 | 2 | N/A | 21 | 78 | N/A | Male | GSE98290 | this paper |
| 29.fastq.gz | CTRL 2 d35 | electrically mature motor neurons | homo sapiens | Patient 3 | CTRL 2 | 1 | N/A | 35 | 78 | N/A | Male | GSE98290 | this paper |
| 30.fastq.gz | CTRL 1 d35 | electrically mature motor neurons | homo sapiens | Patient 4 | CTRL 1 | 1 | N/A | 35 | 64 | N/A | Male | GSE98290 | this paper |
| 31.fastq.gz | CTRL 2 d35 | electrically mature motor neurons | homo sapiens | Patient 3 | CTRL 2 | 2 | N/A | 35 | 78 | N/A | Male | GSE98290 | this paper |

\*different differentiation from same cell line

**Table S3:**
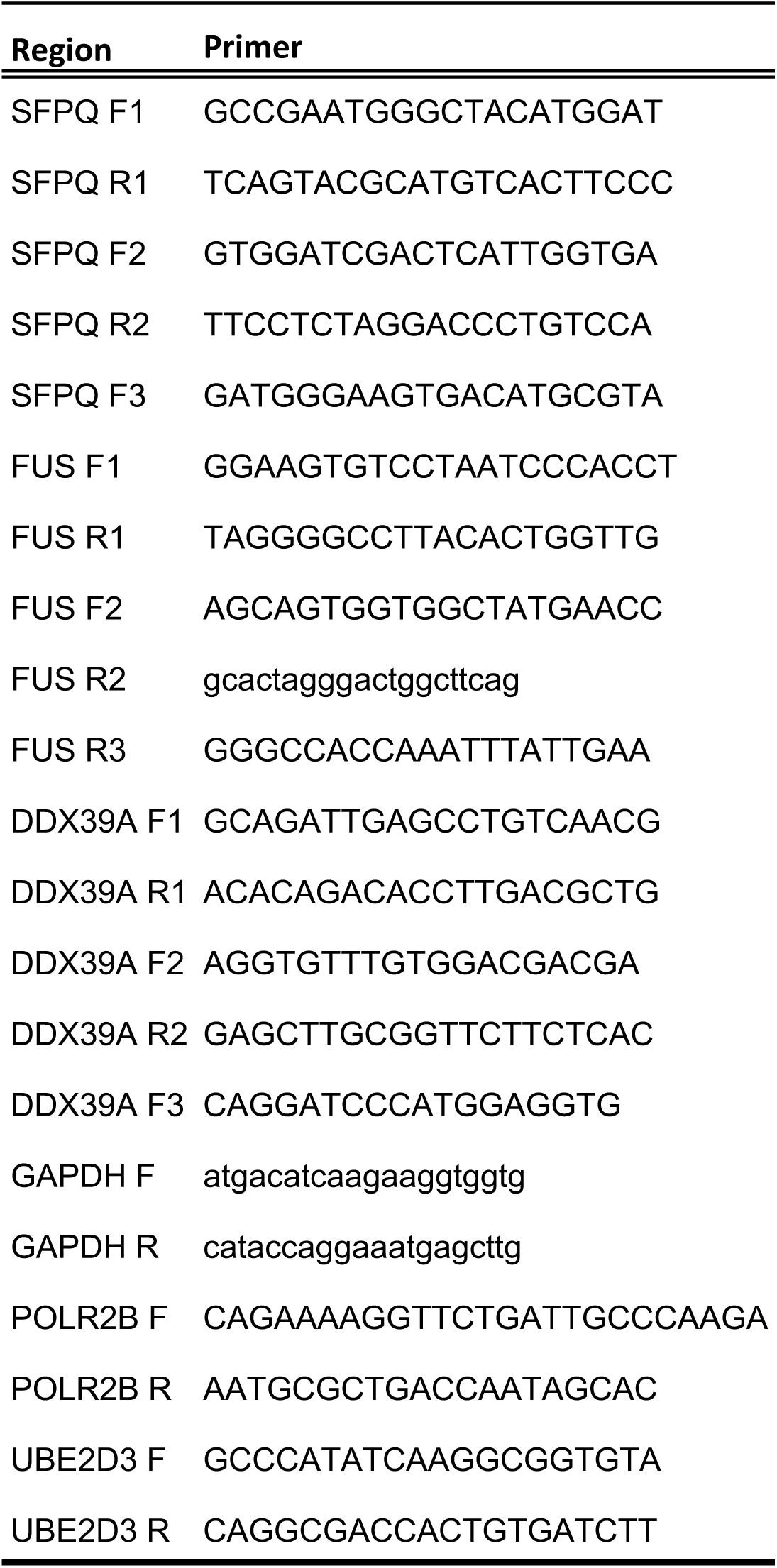
List of primers.

## METHODS

### Contact for reagent and resource sharing

Further information and requests for resources and reagents should be directed to and will be fulfilled by the Lead Contact, Rickie Patani.

### Ethics Statement

Informed consent was obtained from all patients and healthy controls in this study. Experimental protocols were all carried out according to approved regulations and guidelines by UCLH’s National Hospital for Neurology and Neurosurgery and UCL’s Institute of Neurology joint research ethics committee (09/0272).

### Derivation of Human Fibroblasts and iPSC

Dermal fibroblasts were cultured in OptiMEM +10% FCS medium. The following episomal plasmids were transfected for iPSC generation: pCXLE hOct4 shp53, pCXLE hSK, and pCXLE hUL (Addgene), as previously reported (Okita et al., 2011). Details of the lines used in this study are provided in Table S1. Two of the control lines used (control 2 and control 3) are commercially available and were purchased from Coriell (cat. number ND41866*C) and ThermoFisher Scientific (cat. number A18945) respectively.

### Cell Culture

Induced PSCs were maintained on Geltrex (Life Technologies) with Essential 8 Medium media (Life Technologies), and passaged using EDTA (Life Technologies, 0.5mM). All cell cultures were maintained at 37C and 5% carbon dioxide.

### Motor neuron differentiation

Motor neuron (MN) differentiation was carried out using an adapted version of a previously published protocol (Hall et al., 2017). Briefly, iPSCs were first differentiated to neuroepithelium by plating to 100% confluency in chemically defined medium consisting of DMEM/F12 Glutamax, Neurobasal, LGlutamine, N2 supplement, non-essential amino acids, B27 supplement, ß-mercaptoethanol (all from Life Technologies) and insulin (Sigma). Treatment with small molecules from day 0-7 was as follows: 1μM Dorsomorphin (Millipore), 2μM SB431542 (Tocris Bioscience), and 3.3μM CHIR99021 (Miltenyi Biotec). At day 8, the neuroepithelial layer was enzymatically dissociated using dispase (GIBCO, 1 mg/ml), plated onto laminin coated plates and next patterned for 7 days with 0.5μM retinoic acid and 1μM Purmorphamine. At day 14 spinal cord MN precursors were treated with 0.1μM Purmorphamine for a further 4 days before being terminally differentiated in 0.1 μM Compound E (Enzo Life Sciences) to promote cell cycle exit.

### RNA extraction and sequencing

The Promega Maxwell RSC simplyRNA cells kit including DNase treatment, alongside the Maxwell RSC instrument, was used for RNA extractions. For qPCR validations, RNA was extracted using the RNeasy Plus Mini Kit (Qiagen). The nanodrop was used to assess RNA concentration and the 260/280 ratio, and the Agilent bioanalyser was used to assess quality. RNA integrity (RIN) scores were >8 for all samples used in this work. RNAseq libraries were prepared using the Truseq stranded mRNA kit (Illumina) with 1μg input poly(A)+ RNA. The products were then purified and enriched with PCR amplification to create the final cDNA libraries. Libraries were sent for high throughput sequencing, run for 75 cycles on a rapid flow cell, at the Institute of Neurology’s NGS core facility using the Hiseq2500.

### RNA-sequencing data

Single-end stranded RNA-seq reads of 50 bp were obtained from 5 distinct stages of motor neuron differentiation from control and VCP*^mu^* samples (iPSC, and days 7, 14, 21 and 35); samples are listed in Table S2. We also obtained single- and paired-end RNA sequencing reads derived from two independent studies of *in vitro* neural differentiation of hESCs (**GSE20301** (Wu et al., 2010) and **GSE86985** (Yao et al., 2017)); two independent studies on familial forms of ALS either caused by mutant SOD1 (n=5; 2 patient-derived SOD1A4V and 3 isogenic control MN samples where the mutation has been corrected; Hb9 FACS purified MNs, **GSE54409** (Kiskinis et al., 2014a) or FUS (n=6; 3 patient-derived FUS R521G and 3 healthy sibling controls MNs, **GSE77702** (Kapeli et al., 2016); two RNA-seq data from human ESC shared by Miha Modic; Two fetal spMN (**E-MTAB-3871**; NIH Roadmap Epigenomics Mapping Consortium) and adult spMN (laser-captured spMN; **GSE76514** (Nichterwitz et al., 2016)) samples.

### Expression data pre-processing

Single- and paired-reads for each of the study were initially aligned to ribosomal RNA sequences to filter out reads that may come from ribosomal RNA contamination using bowtie2 (-v 0) (Langmead and Salzberg, 2012). The remaining reads were aligned to the human genome (h19) using the splice aware aligner TopHat2 (Kim et al., 2013) with default parameters. All libraries generated in this study had <1% rRNA, <1% mtDNA, >90% strandedness and >70% exonic reads (data not shown).

### Gene quantification and unsupervised characterisation of the data-set

The absolute quantification of the genes was performed using HTSeq count (Anders et al., 2015). Subsequent analysis was performed with the R statistical package version 3.3.1 (2016) and Bioconductor libraries version 3.3 (R Core Team. R: A Language and Environment for Statistical Computing. Vienna, Austria: R Foundation for Statistical Computing; 2013).

Prior to unsupervised clustering analysis of the 31 samples of motor neuron differentiation, we identified reliably expressed genes for each condition (VCP*^mu^* or control at days 0, 7, 14, 21 and 35). For a given sample, the histogram of log2 gene count is generally bimodal, with the modes corresponding to non-expressed and expressed genes. Reliably expressed genes were identified by fitting a two-component Gaussian mixture to the log2 estimated count gene data with R package mclust (Fraley and Raftery). A gene was considered to be reliably expressed in a given condition if the probability of it belonging to the non-expressed class was under 1% in each sample belonging to the condition. 15,989 genes were selected based on their detected expression in at least one of the 10 conditions (i.e. 5 different timepoints of lineage restriction for control and VCP*^mu^*). Next we quantile normalized the columns of the gene count matrix with R package limma (Boldstad et al., 2003). Unsupervised hierarchical clustering of the filtered and normalised gene count matrix was performed with Spearman rank correlation as a distance measure and complete clustering algorithm.

We performed singular value decomposition (SVD) of the expression of the 15,989 genes across the 5 distinct stages of motor neuron differentiation from healthy controls and VCP mutants. We then selected the components maximally capturing variance in gene expression. To visualize the right singular vectors 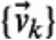, we plotted the expression on the vertical axis as a function of the time corresponding to each sample on the horizontal axis and coloring all samples corresponding to healthy controls gray, and those corresponding to VCP-mutants in magenta. Next we identified genes whose expression profiles correlated (Pearson correlation between individual gene expression profile and right singular vectors) and contributed (projection of each individual gene expression profile onto right singular vectors) most strongly (either positively or negatively) with the expression profile of the singular vectors. In order to identify representative genes for each singular vector, genes were ranked according to both projection and correlation scores. The highest (most positive scores in both projection and correlation) and lowest (most negative scores in both correlation and projection) motifs were selected for each singular vector using K-mean clustering for downstream Gene Ontology enrichment analysis.

### Analysis of alternative polyadenylation from RNA-seq

#### Alternative 3’ UTR identification from RNA-seq

Nucleotide-level stranded coverage was obtained for each of the 31 samples using genomecov from the BEDTools suite (Quinlan, 2014). Next continuously transcribed regions were identified using a sliding window across the genome requiring a minimum coverage of 7 reads in more than 80 positions per window of 100 bp; neighbouring regions separated by low-mappable regions were merged as previously described (Miura et al., 2013). Expressed fragments were then associated with overlapping 3’ UTR using the latest hg19 Ensembl versions v75 (Flicek et al., 2012). Isolated expressed regions that did not overlap with any feature were further associated with the closest 3’ UTR if (1) the closest annotated feature was nothing but a 3’ UTR, (2) if the strand of the expressed region was in line with the strand of the closest 3’ UTR, and (3) if the distance to the 3’ UTR was less than 10’000 kb which is the range of intragenic distance (data not shown). The resulting extended 3’ UTRs were subjected to extensive filtering to exclude potential intragenic transcription, overlapping transcripts, and retained introns as previously described (Miura et al., 2013). We finally intersected the longest 3’ UTR segment annotated with RNA-seq data with a poly(A) site annotation built using reads from 3'-end sequencing libraries in human samples (Gruber et al., 2016) to obtain putative pA within longest 3’ UTR segment of each transcript.

#### Identification of changes in the use of proximal and distal poly(A) sites

We then used the number of reads mapped to -300 nt terminal region of each 3’ UTR isoforms as a proxy for the 3’ UTR isoform expression level. The density of mapped reads in -300 nt terminal region of 3’ UTR isoform is bimodal, with a low-density peak probably corresponding to background transcription i.e. 3’ UTR isoforms of low abundance or 3’ UTR isoforms to which reads were spuriously mapped, and a high-density peak corresponding to expressed 3’ UTR isoforms. In order to identify reliably expressed 3’ UTR isoforms in the study, a two-component Gaussian mixture was fitted to the data using the R package mclust(Fraley and Raftery); an isoform was called reliably expressed if in both replicates had less than 1% chance of belonging to the background category.

In order to identify transcripts that show a marked change in the pA site usage between conditions, we scored the differences in proximal-to-distal poly(A) site usage using the following two scores:

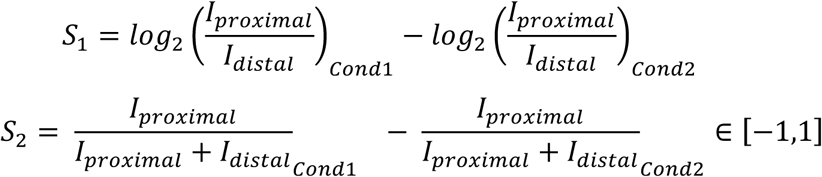

The statistical significance of the changes in proximal-to-distal poly(A) site ratio between two conditions was assessed by Fisher’s exact count test using summed-up raw read counts of promoter-proximal versus promoter-distal 3’ UTR isoforms originating either conditions. We adjusted the P-Value controlling for False Discovery Rate (FDR) of 0.01. We restricted our analysis on Ensembl transcripts containing at least two 3’ UTRs generated by tandem polyadenylation expressed in conditions of interest. Proximal shifts were then selected when *S*_1_ ≤ -1, *S*_2_ ≤ -15% and FDR<0.01; distal shift were selected when selected when *S*_1_ ≥ 1, *S*_2_ ≥ 15% and FDR<0.01.

### Splicing analysis

The identification of all classes of alternative splicing (AS) events in motor neuron differentiation was performed with the RNA-seq pipeline *vast-tools* (Irimia et al., 2014). For an AS event to be considered differentially regulated between two conditions, we required a minimum average ΔPSI (between the paired replicates) of at least 10% and that the transcript targeted by the splicing event in question to be reliably expressed in all samples from the conditions compared i.e enough read coverage in all samples of interest. Intron retention (IR) focussed analysis has next been performed on 167 IR events for which a percentage of IR has been calculated as the fraction of intron mapping reads to the average number of reads mapping to the adjacent 5’ and 3’ exons normalised to the length of the respective intron and exons. A Fisher count test P-value has been obtained when testing for differential IR between conditions.

### Gene ontology enrichment analysis

GO enrichment analysis was performed using classic Fisher test with topGO Bioconductor package (Alexa and Rahnenfuhrer, 2016). Only GO terms containing at least 10 annotated genes were considered. A p-value of 0.05 was used as the level of significance. On the figures, top significant GO terms were manually selected by removing redundant GO terms and terms which contain fewer than 5 significant genes.

### Network analysis

Experimental protein-protein interaction information has been retrieved from the STRING data-base (Szklarczyk et al., 2011) and visualised in Cytoscape (Shannon et al., 2003).

### Reverse transcription, qPCR and intron retention validation

Reverse transcription was performed using the Revert Aid First Strand cDNA Synthesis Kit (ThermoFisher Scientific) using 1μg of total RNA and random hexamers. qPCR was performed using the Power SYBR Green Master Mix (ThermoFisher Scientific) and the Agilent Mx3000P QPCR System. Primers used are listed in Table S3. For each intron retention validation to screen three primers pairs were used: (1) primer pair F1 R1 (intron spanning, across exon-exon junction) were used to to analyse gene expression levels; (2) primer pair F2 R2 (one primer on an exon flanking the intron to be analysed, the other on the intron) was used to assess the levels of intron retention; (3) primer pair F3 R2 (both primers on the exons flanking the intron of interest, if possible designed across the exon-exon junction) was used to measure levels of the spliced transcript. RT-minus samples were used as negative controls. Levels of intron retention (primer pair F2R2) were normalised over the expression level of each individual gene (primer pair F1R1). Gene expression levels were measured using the *ddCt* method using three housekeeping genes (GAPDH, POLR2B and UBE2D3).

### Cell Cycle analysis

Cell cycle analysis was performed by flow cytometry according to standard protocols. Briefly, cells were dissociated using either Accutase or Trypsin (Life Technologies), washed and fixed in suspension using 70% cold Ethanol. Cells were stained using propidium iodide (PI, 50μg/ml, Sigma Aldrich) in the presence of RNase A (10μg/ml, Sigma Aldrich). Cells were analysed using a BD FACS Calibur (BD Bioscience). Doublets were excluded from analysis and 10,000 events were collected in the single cell gate per sample. Cell cycle data was analysed using the Multicycle module of FCS Express 6 (De Novo Software).

## REFERENCES

Alexa, A., and Rahnenfuhrer, J. (2016). topGO: Enrichment Analysis for Gene Ontology.

Anders, S., Pyl, P.T., and Huber, W. (2015). HTSeq—a Python framework to work with high-throughput sequencing data. Bioinformatics 31, 166–169.

Arnold, E.S., Ling, S.-C., Huelga, S.C., Lagier-Tourenne, C., Polymenidou, M., Ditsworth, D., Kordasiewicz, H.B., McAlonis-Downes, M., Platoshyn, O., Parone, P.A., et al. (2013). ALS-linked TDP-43 mutations produce aberrant RNA splicing and adult-onset motor neuron disease without aggregation or loss of nuclear TDP-43. Proc. Natl. Acad. Sci. U. S. A. 110, E736–E745.

Boldstad, B.M., Irizarry, R.A., Astrand, M., and Speed, T.P. (2003). A Comparison of Normalization Methods for High Density Oligonucleotide Array Data Based on Bias and Variance. Bioinformatics 19, 185–193.

Bond, C.S., and Fox, A.H. (2009). Paraspeckles: nuclear bodies built on long noncoding RNA. J. Cell Biol. 186, 637–644.

Braunschweig, U., Barbosa-Morais, N.L., Pan, Q., Nachman, E.N., Alipanahi, B., Gonatopoulos-Pournatzis, T., Frey, B., Irimia, M., and Blencowe, B.J. (2014). Widespread intron retention in mammals functionally tunes transcriptomes. Genome Res. 24, 1774–1786.

Buckley, P.T., Lee, M.T., Sul, J.-Y., Miyashiro, K.Y., Bell, T.J., Fisher, S.A., Kim, J., and Eberwine, J. (2011). Cytoplasmic intron sequence-retaining transcripts can be dendritically targeted via ID element retrotransposons. Neuron 69, 877–884.

Chen, L.-L., and Carmichael, G.G. (2009). Altered nuclear retention of mRNAs containing inverted repeats in human embryonic stem cells: functional role of a nuclear noncoding RNA. Mol. Cell 35, 467–478.

Clark, J.A., Southam, K.A., Blizzard, C.A., King, A.E., and Dickson, T.C. (2016). Axonal degeneration, distal collateral branching and neuromuscular junction architecture alterations occur prior to symptom onset in the SOD1(G93A) mouse model of amyotrophic lateral sclerosis. J. Chem. Neuroanat. 76, 35–47.

Cosker, K.E., Fenstermacher, S.J., Pazyra-Murphy, M.F., Elliott, H.L., and Segal, R.A. (2016). The RNA-binding protein SFPQ orchestrates an RNA regulon to promote axon viability. Nat. Neurosci. 19, 690–696.

Danckwardt, S., Kaufmann, I., Gentzel, M., Foerstner, K.U., Gantzert, A.-S., Gehring, N.H., Neu-Yilik, G., Bork, P., Keller, W., Wilm, M., et al. (2007). Splicing factors stimulate polyadenylation via USEs at non-canonical 3’ end formation signals. EMBO J. 26, 2658–2669.

Derti, A., Garrett-Engele, P., MacIsaac, K.D., Stevens, R.C., Sriram, S., Chen, R., Rohl, C.A., Johnson, J.M., and Babak, T. (2012). A quantitative atlas of polyadenylation in five mammals. Genome Res. 22, 1173–1183.

Dillman, A.A., Hauser, D.N., Raphael Gibbs, J., Nalls, M.A., McCoy, M.K., Rudenko, I.N., Galter, D., and Cookson, M.R. (2013). mRNA expression, splicing and editing in the embryonic and adult mouse cerebral cortex. Nat. Neurosci. 16, 499–506.

Dong, X., Sweet, J., Challis, J.R.G., Brown, T., and Lye, S.J. (2007). Transcriptional activity of androgen receptor is modulated by two RNA splicing factors, PSF and p54nrb. Mol. Cell. Biol. 27, 4863–4875.

Fabian, M.R., Sonenberg, N., and Filipowicz, W. (2010). Regulation of mRNA translation and stability by microRNAs. Annu. Rev. Biochem. 79, 351–379.

Flicek, P., Ahmed, I., Amode, M.R., Barrell, D., Beal, K., Brent, S., Carvalho-Silva, D., Clapham, P., Coates, G., Fairley, S., et al. (2012). Ensembl 2013. Nucleic Acids Res. gks1236.

Fraley, C., and Raftery, A.E. mclust Version 4 for R: Normal Mixture Modeling for Model-Based Clustering, Classification, and Density Estimation.

Gabut, M., Chaudhry, S., and Blencowe, B.J. (2008). SnapShot: The splicing regulatory machinery. Cell 133, 192.e1.

Gruber, A.J., Schmidt, R., Gruber, A.R., Martin, G., Ghosh, S., Belmadani, M., Keller, W., and Zavolan, M. (2016). A comprehensive analysis of 3’ end sequencing data sets reveals novel polyadenylation signals and the repressive role of heterogeneous ribonucleoprotein C on cleavage and polyadenylation. Genome Res. 26, 1145–1159.

Gupta, I., Clauder-Münster, S., Klaus, B., Järvelin, A.I., Aiyar, R.S., Benes, V., Wilkening, S., Huber, W., Pelechano, V., and Steinmetz, L.M. (2014). Alternative polyadenylation diversifies post-transcriptional regulation by selective RNA-protein interactions. Mol. Syst. Biol. 10, 719.

Hall, C.E., Yao, Z., Choi, M., Tyzack, G.E., Serio, A., Luisier, R., Harley, J., Preza, E., Arber, C., Crisp, S.J., et al. (2017). Progressive Motor Neuron Pathology and the Role of Astrocytes in a Human Stem Cell Model of VCP-Related ALS. Cell Rep. 19, 1739–1749.

Hall-Pogar, T., Liang, S., Hague, L.K., and Lutz, C.S. (2007). Specific trans-acting proteins interact with auxiliary RNA polyadenylation elements in the COX-2 3'-UTR. RNA 13, 1103–1115.

Heyn, P., Kalinka, A.T., Tomancak, P., and Neugebauer, K.M. (2015). Introns and gene expression: cellular constraints, transcriptional regulation, and evolutionary consequences. Bioessays 37, 148–154.

Hirose, T., Virnicchi, G., Tanigawa, A., Naganuma, T., Li, R., Kimura, H., Yokoi, T., Nakagawa, S., Bénard, M., Fox, A.H., et al. (2014). NEAT1 long noncoding RNA regulates transcription via protein sequestration within subnuclear bodies. Mol. Biol. Cell 25, 169–183.

Ho, R., Sances, S., Gowing, G., Amoroso, M.W., O'Rourke, J.G., Sahabian, A., Wichterle, H., Baloh, R.H., Sareen, D., and Svendsen, C.N. (2016). ALS disrupts spinal motor neuron maturation and aging pathways within gene co-expression networks. Nat. Neurosci. 19, 1256–1267.

Irimia, M., Weatheritt, R.J., Ellis, J.D., Parikshak, N.N., Gonatopoulos-Pournatzis, T., Babor, M., Quesnel-Vallières, M., Tapial, J., Raj, B., O'Hanlon, D., et al. (2014). A highly conserved program of neuronal microexons is misregulated in autistic brains. Cell 159, 1511–1523.

Ishigaki, S., Fujioka, Y., Okada, Y., Riku, Y., Udagawa, T., Honda, D., Yokoi, S., Endo, K., Ikenaka, K., Takagi, S., et al. (2017). Altered Tau Isoform Ratio Caused by Loss of FUS and SFPQ Function Leads to FTLD-like Phenotypes. Cell Rep. 18, 1118–1131.

Ji, Z., Lee, J.Y., Pan, Z., Jiang, B., and Tian, B. (2009). Progressive lengthening of 3’ untranslated regions of mRNAs by alternative polyadenylation during mouse embryonic development. Proceedings of the National Academy of Sciences 106, 7028–7033.

Johnson, J.O., Mandrioli, J., Benatar, M., Abramzon, Y., Van Deerlin, V.M., Trojanowski, J.Q., Gibbs, J.R., Brunetti, M., Gronka, S., Wuu, J., et al. (2010). Exome sequencing reveals VCP mutations as a cause of familial ALS. Neuron 68, 857–864.

Kapeli, K., Pratt, G.A., Vu, A.Q., Hutt, K.R., Martinez, F.J., Sundararaman, B., Batra, R., Freese, P., Lambert, N.J., Huelga, S.C., et al. (2016). Distinct and shared functions of ALS-associated proteins TDP-43, FUS and TAF15 revealed by multisystem analyses. Nat. Commun. 7, 12143.

Kapeli, K., Martinez, F.J., and Yeo, G.W. (2017). Genetic mutations in RNA-binding proteins and their roles in ALS. Hum. Genet.

Ke, Y.D., Ke, Y., Dramiga, J., Schütz, U., Kril, J.J., Ittner, L.M., Schröder, H., and Götz, J. (2012). Tau-mediated nuclear depletion and cytoplasmic accumulation of SFPQ in Alzheimer's and Pick's disease. PLoS One 7, e35678.

Kerschbamer, E., and Biagioli, M. (2015). Huntington's Disease as Neurodevelopmental Disorder: Altered Chromatin Regulation, Coding, and Non-Coding RNA Transcription. Front. Neurosci. 9, 509.

Kim, D., Pertea, G., Trapnell, C., Pimentel, H., Kelley, R., and Salzberg, S.L. (2013). TopHat2: accurate alignment of transcriptomes in the presence of insertions, deletions and gene fusions. Genome Biol. 14, R36.

Kiskinis, E., Sandoe, J., Williams, L.A., Boulting, G.L., Moccia, R., Wainger, B.J., Han, S., Peng, T., Thams, S., Mikkilineni, S., et al. (2014a). Pathways disrupted in human ALS motor neurons identified through genetic correction of mutant SOD1. Cell Stem Cell 14, 781–795.

Kiskinis, E., Sandoe, J., Williams, L.A., Boulting, G.L., Moccia, R., Wainger, B.J., Han, S., Peng, T., Thams, S., Mikkilineni, S., et al. (2014b). Pathways disrupted in human ALS motor neurons identified through genetic correction of mutant SOD1. Cell Stem Cell 14, 781–795.

Langmead, B., and Salzberg, S.L. (2012). Fast gapped-read alignment with Bowtie 2. Nat. Methods 9, 357–359.

Licatalosi, D.D., Mele, A., Fak, J.J., Ule, J., Kayikci, M., Chi, S.W., Clark, T.A., Schweitzer, A.C., Blume, J.E., Wang, X., et al. (2008). HITS-CLIP yields genome-wide insights into brain alternative RNA processing. Nature 456, 464–469.

Lourenco, G.F., Janitz, M., Huang, Y., and Halliday, G.M. (2015). Long noncoding RNAs in TDP-43 and FUS/TLS-related frontotemporal lobar degeneration (FTLD). Neurobiol. Dis. 82, 445–454.

Luisier, R., Unterberger, E.B., Goodman, J.I., Schwarz, M., Moggs, J., Terranova, R., and van Nimwegen, E. (2014). Computational modeling identifies key gene regulatory interactions underlying phenobarbital-mediated tumor promotion. Nucleic Acids Res. gkt1415.

Mansfield, K.D., and Keene, J.D. (2011). Neuron-specific ELAV/Hu proteins suppress HuR mRNA during neuronal differentiation by alternative polyadenylation. Nucleic Acids Res. gkr1114.

Masuda, A., Takeda, J.-I., and Ohno, K. (2016). FUS-mediated regulation of alternative RNA processing in neurons: insights from global transcriptome analysis. Wiley Interdiscip. Rev. RNA.

Mauger, O., Lemoine, F., and Scheiffele, P. (2016). Targeted Intron Retention and Excision for Rapid Gene Regulation in Response to Neuronal Activity. Neuron 92, 1266–1278.

Miura, P., Shenker, S., Andreu-Agullo, C., Westholm, J.O., and Lai, E.C. (2013). Widespread and extensive lengthening of 3' UTRs in the mammalian brain. Genome Res. 23, 812–825.

Naganuma, T., Nakagawa, S., Tanigawa, A., Sasaki, Y.F., Goshima, N., and Hirose, T. (2012). Alternative 3’-end processing of long noncoding RNA initiates construction of nuclear paraspeckles. EMBO J. 31, 4020–4034.

Neumann, M., Sampathu, D.M., Kwong, L.K., Truax, A.C., Micsenyi, M.C., Chou, T.T., Bruce, J., Schuck, T., Grossman, M., Clark, C.M., et al. (2006). Ubiquitinated TDP-43 in frontotemporal lobar degeneration and amyotrophic lateral sclerosis. Science 314, 130–133.

Nichterwitz, S., Chen, G., Aguila Benitez, J., Yilmaz, M., Storvall, H., Cao, M., Sandberg, R., Deng, Q., and Hedlund, E. (2016). Laser capture microscopy coupled with Smart-seq2 for precise spatial transcriptomic profiling. Nat. Commun. 7, 12139.

Nilsen, T.W., and Graveley, B.R. (2010). Expansion of the eukaryotic proteome by alternative splicing. Nature 463, 457–463.

Nishimoto, Y., Nakagawa, S., Hirose, T., Okano, H.J., Takao, M., Shibata, S., Suyama, S., Kuwako, K.-I., Imai, T., Murayama, S., et al. (2013). The long non-coding RNA nuclear-enriched abundant transcript 1_2 induces paraspeckle formation in the motor neuron during the early phase of amyotrophic lateral sclerosis. Mol. Brain 6, 31.

Okita, K., Matsumura, Y., Sato, Y., Okada, A., Morizane, A., Okamoto, S., Hong, H., Nakagawa, M., Tanabe, K., Tezuka, K.-I., et al. (2011). A more efficient method to generate integration-free human iPS cells. Nat. Methods 8, 409–412.

Patton, J.G., Porro, E.B., Galceran, J., Tempst, P., and Nadal-Ginard, B. (1993). Cloning and characterization of PSF, a novel pre-mRNA splicing factor. Genes Dev. 7, 393–406.

Qiu, H., Lee, S., Shang, Y., Wang, W.-Y., Au, K.F., Kamiya, S., Barmada, S.J., Finkbeiner, S., Lui, H., Carlton, C.E., et al. (2014). ALS-associated mutation FUSR521C causes DNA damage and RNA splicing defects. J. Clin. Invest. 124, 981–999.

Quinlan, A.R. (2014). BEDTools: The Swiss-Army Tool for Genome Feature Analysis. Curr. Protoc. Bioinformatics 11–12.

Raj, B., Irimia, M., Braunschweig, U., Sterne-Weiler, T., O'Hanlon, D., Lin, Z.-Y., Chen, G.I., Easton, L.E., Ule, J., Gingras, A.-C., et al. (2014). A global regulatory mechanism for activating an exon network required for neurogenesis. Mol. Cell 56, 90–103.

Sasaki, Y.T.F., Ideue, T., Sano, M., Mituyama, T., and Hirose, T. (2009). MENepsilon/beta noncoding RNAs are essential for structural integrity of nuclear paraspeckles. Proc. Natl. Acad. Sci. U. S. A. 106, 2525–2530.

Schultz, M.D., He, Y., Whitaker, J.W., Hariharan, M., Mukamel, E.A., Leung, D., Rajagopal, N., Nery, J.R., Urich, M.A., Chen, H., et al. (2015). Human body epigenome maps reveal noncanonical DNA methylation variation. Nature 523, 212–216.

Shannon, P., Markiel, A., Ozier, O., Baliga, N.S., Wang, J.T., Ramage, D., Amin, N., Schwikowski, B., and Ideker, T. (2003). Cytoscape: a software environment for integrated models of biomolecular interaction networks. Genome Res. 13, 2498–2504.

Szklarczyk, D., Franceschini, A., Kuhn, M., Simonovic, M., Roth, A., Minguez, P., Doerks, T., Stark, M., Muller, J., Bork, P., et al. (2011). The STRING database in 2011: functional interaction networks of proteins, globally integrated and scored. Nucleic Acids Res. 39, D561–D568.

Szklarczyk, D., Morris, J.H., Cook, H., Kuhn, M., Wyder, S., Simonovic, M., Santos, A., Doncheva, N.T., Roth, A., Bork, P., et al. (2017). The STRING database in 2017: quality-controlled protein-protein association networks, made broadly accessible. Nucleic Acids Res. 45, D362–D368.

Thomas-Jinu, S., Gordon, P.M., Fielding, T., Taylor, R., Smith, B.N., Snowden, V., Blanc, E., Vance, C., Topp, S., Wong, C.-H., et al. (2017). Non-nuclear Pool of Splicing Factor SFPQ Regulates Axonal Transcripts Required for Normal Motor Development. Neuron 94, 931.

Tian, B., and Manley, J.L. (2017). Alternative polyadenylation of mRNA precursors. Nat. Rev. Mol. Cell Biol. 18, 18–30.

Vance, C., Rogelj, B., Hortobágyi, T., De Vos, K.J., Nishimura, A.L., Sreedharan, J., Hu, X., Smith, B., Ruddy, D., Wright, P., et al. (2009). Mutations in FUS, an RNA processing protein, cause familial amyotrophic lateral sclerosis type 6. Science 323, 1208–1211.

Wang, L., and Yi, R. (2014). 3’UTRs take a long shot in the brain. Bioessays 36, 39–45.

Ward, L.D., and Kellis, M. (2012). Interpreting noncoding genetic variation in complex traits and human disease. Nat. Biotechnol. 30, 1095–1106.

Will, T.J., Tushev, G., Kochen, L., Nassim-Assir, B., Cajigas, I.J., Schuman, E.M., and Others (2013). Deep sequencing and high-resolution imaging reveal compartment-specific localization of Bdnf mRNA in hippocampal neurons. Sci. Signal. 6, rs16.

Wu, J.Q., Habegger, L., Noisa, P., Szekely, A., Qiu, C., Hutchison, S., Raha, D., Egholm, M., Lin, H., Weissman, S., et al. (2010). Dynamic transcriptomes during neural differentiation of human embryonic stem cells revealed by short, long, and paired-end sequencing. Proc. Natl. Acad. Sci. U. S. A. 107, 5254–5259.

Yan, Q., Weyn-Vanhentenryck, S.M., Wu, J., Sloan, S.A., Zhang, Y., Chen, K., Wu, J.Q., Barres, B.A., and Zhang, C. (2015). Systematic discovery of regulated and conserved alternative exons in the mammalian brain reveals NMD modulating chromatin regulators. Proceedings of the National Academy of Sciences 112, 3445–3450.

Yao, Z., Mich, J.K., Ku, S., Menon, V., Krostag, A.-R., Martinez, R.A., Furchtgott, L., Mulholland, H., Bort, S., Fuqua, M.A., et al. (2017). A Single-Cell Roadmap of Lineage Bifurcation in Human ESC Models of Embryonic Brain Development. Cell Stem Cell 20, 120–134.

Yap, K., Lim, Z.Q., Khandelia, P., Friedman, B., and Makeyev, E.V. (2012). Coordinated regulation of neuronal mRNA steady-state levels through developmentally controlled intron retention. Genes Dev. 26, 1209–1223.

Yap, K., Xiao, Y., Friedman, B.A., Je, H.S., and Makeyev, E.V. (2016). Polarizing the Neuron through Sustained Co-expression of Alternatively Spliced Isoforms. Cell Rep. 15, 1316–1328.

Yarosh, C.A., Iacona, J.R., Lutz, C.S., and Lynch, K.W. (2015). PSF: nuclear busy-body or nuclear facilitator? Wiley Interdiscip. Rev. RNA 6, 351–367.

Zhang, H., Lee, J.Y., and Tian, B. (2005). Biased alternative polyadenylation in human tissues. Genome Biol. 6, R100.

Zhang, X., Chen, M.H., Wu, X., Kodani, A., Fan, J., Doan, R., Ozawa, M., Ma, J., Yoshida, N., Reiter, J.F., et al. (2016). Cell-Type-Specific Alternative Splicing Governs Cell Fate in the Developing Cerebral Cortex. Cell 166, 1147–1162.e15.

